# Parallel Labeled-Line Organization of Sympathetic Outflow for Selective Organ Regulation in Mice

**DOI:** 10.1101/2024.04.04.588164

**Authors:** Yukiko Harima, Masafumi Tsurutani, Shuntaro Uchida, Kengo Inada, Mitsue Hagihara, Satsuki Irie, Mayo Shigeta, Takaya Abe, Yukiko U. Inoue, Takayoshi Inoue, Kazunari Miyamichi

## Abstract

The sympathetic nervous system is vital in maintaining homeostasis and responding to environmental changes^1–3^. This regulation is coordinated by the spinal sympathetic preganglionic neurons (SPNs), which influence various organs both through neuronal pathways via postganglionic neurons and through endocrine processes by innervating the adrenal gland. Despite decades of research supporting the concept of selective control within this system^1,4–9^, the neural circuit organization responsible for the specificity of sympathetic outflow remains poorly understood. Notably, classical anatomical studies in rats have not revealed a definitive molecular code governing SPNs, nor have they confirmed the existence of SPNs strictly corresponding to specific output targets^1,6,10,11^. To reconcile this discrepancy, we aim to integrate recent transcriptome data of SPNs^12,13^ in mice with viral-genetic toolkits^14^ to map axonal projections and manipulate the functions of SPNs governing the gastrointestinal tract and adrenal gland. Here, we have identified two subtypes of SPNs in the lower thoracic spinal cord, defined at the molecular level, exhibiting non-overlapping patterns of innervation. Chemogenetic manipulations on these distinct SPN subtypes revealed selective impacts on the digestive functions in the gastrointestinal tracts or glucose metabolism mediated by the adrenal gland, respectively. This molecularly delineated parallel labeled-line organization in sympathetic outflows presents a potential avenue for selectively manipulating organ functions.

---

Recent advancements in single-cell transcriptome (TC) technologies have expanded neuronal cell type classification^15^, raising general questions regarding whether a given TC type projects exclusively to a specific set of target regions (labeled-line model, Fig. 1a_1_) or if multiple TC types within a given brain region share common output targets (divergent output model, Fig. 1a_2_). While recent studies have exemplified varying degrees of output specificity among TC types across multiple central brain regions^16,17^, this issue within the efferent pathways in the spinal cord (SC) remains unexplored, despite its significant relevance to the regulation of various organs in the body. In this context, here we focus on the SPNs that target either the celiac/superior mesenteric ganglia (CG/SMG) or the adrenal medulla (AM), two crucial components of the fight-or-flight response. Upon encountering stressors, the sympathetic nervous system provides energy supply, partly via the function of the AM^18^, while concurrently inhibiting the digestive functions in the gastrointestinal (GI) tract via the post-ganglionic neurons (postGNs) located in the CG/SMG^19^. The SPNs regulating these targets are located within the same lower thoracic SC^1^, which renders these SPNs an ideal model for studying their molecular, anatomical, and functional specificities. Notably, animals can differentially utilize these two output pathways; AM-projecting SPNs become more active in response to low plasma glucose levels^6^, and AM chromaffin cells are selectively activated by a brain-infused hormone^7^. Despite these observations supporting the presence of distinct output specificity, classical anatomical studies in rats neither reveal a definitive molecular code for SPNs nor corroborate the labeled line organization^1,6,10,11^. This motivated us to utilize recent transcriptome data of SPNs^12,13^ in mice and viral-genetic toolkits^14^ to map axonal projections and manipulate the functions of SPNs targeting the CG/SMG or the AM.

**Fig. 1:**
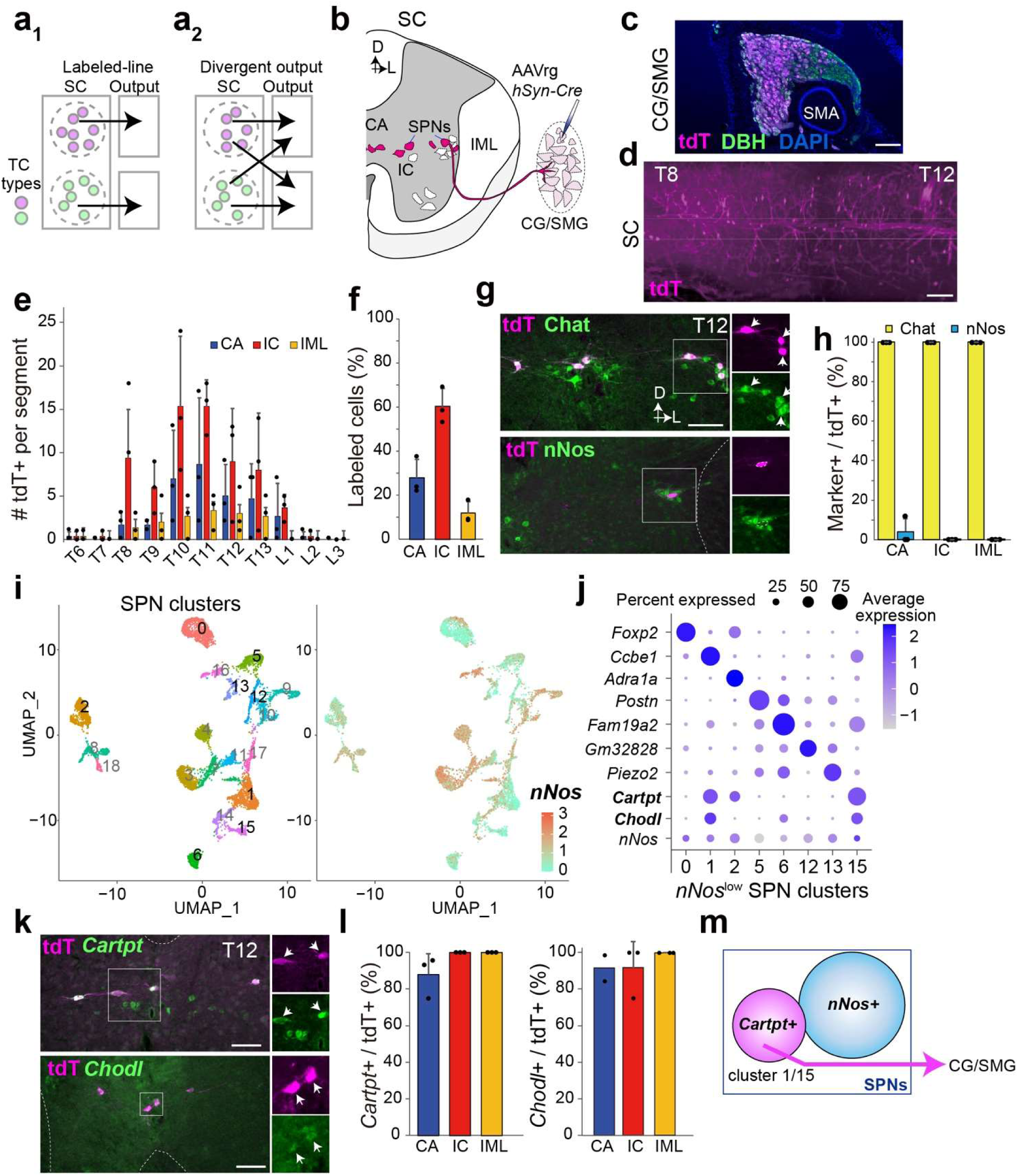
Molecular identities of CG/SMG-projecting SPNs. **a**, Schematic of two extreme projection patterns for two distinct TC types. **b**, Schematic of retrograde tracing from the CG/SMG. CA, central autonomic area; IC, intercalated nucleus; IML, intermediolateral cell column; D, dorsal; L, lateral. **c**, Horizontal view of the CG/SMG of *Ai9* mice injected with AAVrg-*hSyn-Cre*. tdT, tdTomato from *Ai9* mice. DBH, dopamine-β-hydroxylase, which is a functional marker of the sympathetic postGNs. SMA, superior mesenteric artery. Scale bar, 200 μm. **d**, Light-sheet microscopic view of tissue-cleared SC displaying retrogradely labeled tdT+ cells (magenta). Scale bar, 500 μm. T, thoracic. **e**, Number of tdT+ cells in the CA (blue), IC (red), and IML (yellow) for each SC segment. N = 3 mice. L, lumbar. **f**, Percentage of labeled cells in the CA (blue), IC (red), and IML (yellow). N = 3 mice. **g**, Coronal sections of the T12 SC depicting retrogradely labeled tdT+ cells (magenta) and Chat (top) or nNos (bottom) immunostaining (green). Magnified images within white boxes are displayed on the right of each section. The arrows indicate cells that are double-positive for tdT and Chat. Scale bars, 100 μm. **h**, Percentage of Chat+ or nNos+ cells among tdT+ cells in the CA, IC, and IML. N = 3 mice. Data are presented as mean ± standard deviation. **i**, UMAP representation of 19 *Chat*+ SPN clusters based on the published transcriptome data^12^ (left) and a colored heat map showing log-normalized expression of the *nNos* gene within SPN clusters (right). **j**, Identification of enriched differentially expressed genes and expression of *nNos* among SPNs for *nNos*^low^ clusters. The dot size reflects the percentage of cells within each cluster expressing the respective marker gene, while the intensity of the blue color indicates the log-normalized expression level. **k**, Coronal sections of the T12 SC displaying retrogradely labeled tdT+ cells (magenta) from the CG/SMG and *Cartpt* (top) or *Chodl* (bottom) mRNA expression (green). Magnified and channel-separated images within the white box are presented on the right. Arrows indicate dual-labeled cells. Scale bars, 100 μm. **l**, Percentage of *Cartpt*+ cells (left) or *Chodl*+ cells (right) among tdT+ cells. N = 2–3 mice. **m**, Schematics summary of the data. Data are presented as mean ± standard deviation. For more data, see Extended Data Fig. 1 and Supplementary Movies 1 and 2.

## *Cartpt*+ SPNs selectively project to the CG/SMG

To unravel the molecular characteristics of SPNs projecting to the CG/SMG, we developed selective adeno-associated virus (AAV) injection procedures into CG/SMG in mice (Extended Data Fig. 1, Supplementary Movie 1). We then injected retrogradely transducible AAVrg *hSyn-Cre* into the CG/SMG of *Ai9* (Cre reporter tdTomato line^20^) mice to visualize SPNs in the SC (Fig. 1b, c). Under light sheet microscopy after tissue clearance^21^, tdTomato-expressing (+) cells retrogradely labeled from the CG/SMG were predominantly found in the 8th to 13th thoracic SC (T8–T13) (Fig. 1d, Supplementary Movie 2). Histological analysis on coronal SC sections confirmed this distribution and also revealed that over 80% of tdTomato+ cells were located in the central autonomic area (CA) and intercalated nucleus (IC), contrary to the prevailing view that the majority of SPNs are located in the intermediolateral cell column (IML)^22^ (Fig. 1e, f). While tdTomato+ cells were nearly 100% positive for choline acetyltransferase (Chat), a functional marker of cholinergic SPNs (Fig. 1g, h), there was no overlap between tdTomato+ cells and those labeled with neuronal nitric oxide synthase, nNos, a conventional marker for SPNs^23,24^ (Fig. 1g, h).

The detailed molecular profiles of SPNs with low or no expression of nNos have not been adequately addressed in previous studies. To characterize them, we conducted a reanalysis of existing single-nucleus RNA sequencing (snRNAseq) data by Blum *et al.*^12^ (see Methods for details). We defined *nNos*^low^ SPN clusters as those in which 50% or fewer cells express *nNos*, with average expression levels (in arbitrary units) lower than zero. This led to the identification of 8 *nNos*^low^ SPN clusters (Fig. 1i, j). Histochemical analyses using selective marker genes for these *nNos*^low^ clusters revealed that tdTomato+ cells retrogradely labeled from the CG/SMG almost entirely co-localized with those expressing *Cartpt* (Fig. 1k, l), a dominant marker of clusters 1 and 15 with relatively weak expression in clusters 2 (Fig. 1j). We also observed that the vast majority of tdTomato+ cells expressed the *Chondrolectin gene* (*Chodl*), marking clusters 1, and 15 (Fig. 1j–l), suggesting that the tdTomato+ cells belong to SPN clusters 1 and 15. Collectively, these data indicate that SPNs retrogradely labeled from the CG/SMG form molecularly defined distinct TC subtypes marked by *Cartpt* (Fig. 1m).

To analyze the axonal projection targets of *Cartpt*+ SPNs, we introduced an AAV expressing *Cre*-dependent mCherry into the T8–T12 of mice that expressed *Cre* under the control of the *Cartpt*, with pan-SPN driver *Chat-Cre* mice^25^ served as a positive control (Fig. 2a, b). *Cartpt-Cre* mice were generated through the CRISPR/Cas9-based method^26^ (Extended Data Fig. 2). At 2–3 weeks post-viral injection, we confirmed that the majority of mCherry+ cells were Chat+ or *Cartpt*+, respectively, demonstrating high specificity (Fig. 2c). We analyzed their axons in the sections of CG/SMG, AM, and the sympathetic trunk (ST) (Fig. 2d). While *Chat*+ pan-SPN axons were closely co-localized with the dopamine-β-hydroxylase (DBH)+ postGNs or adrenal chromaffin cells in these structures, we exclusively detected mCherry+ axons of *Cartpt*+ SPNs in only the CG/SMG, with no presence in the AM or ST (Fig. 2d).

**Fig. 2:**
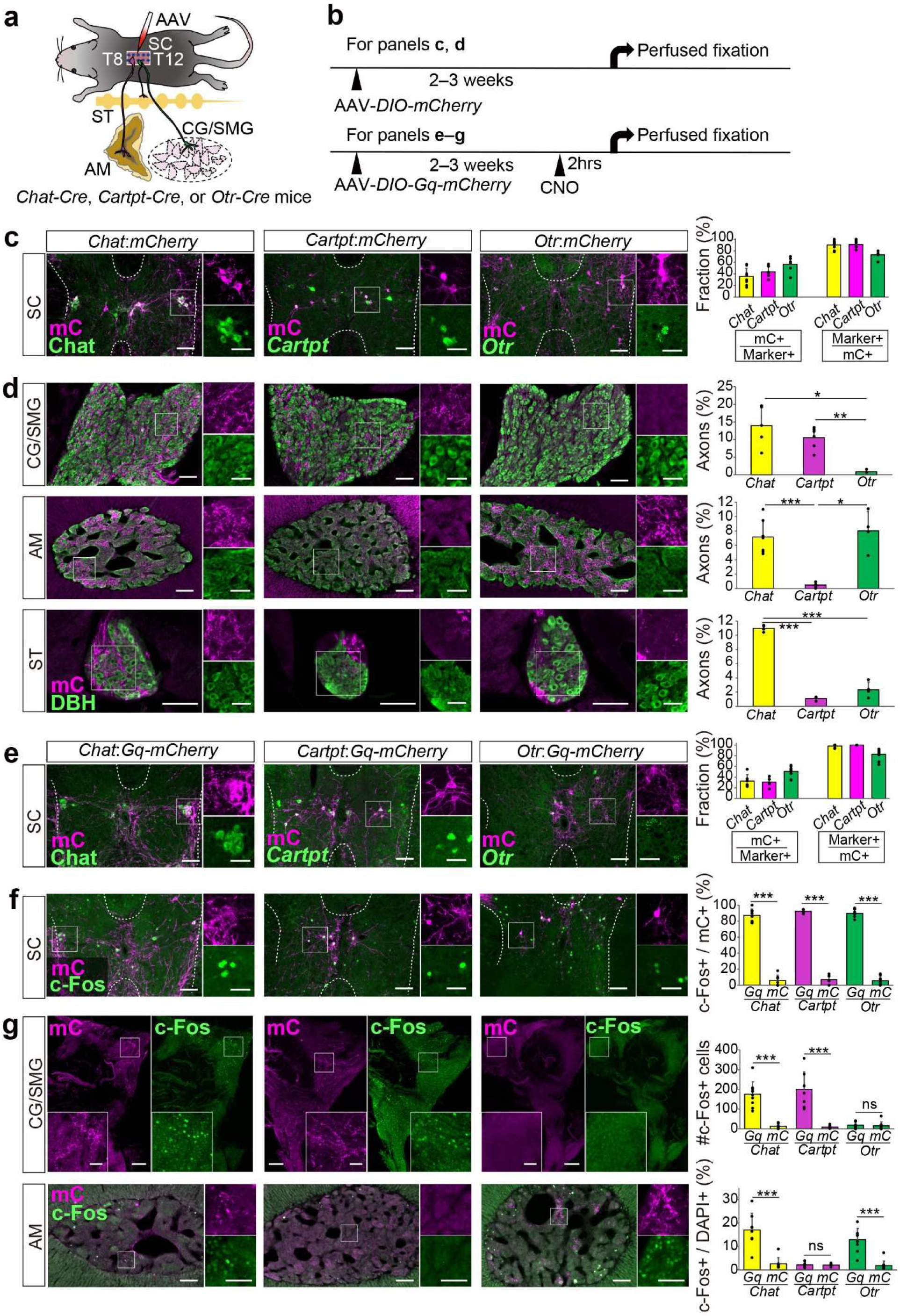
Distinct projection targets of *Cartpt*+ and *Otr*+ SPNs. **a, b**, Schematics illustrating the experimental design (**a**) and time line (**b**). AM, adrenal medulla. ST, sympathetic trunk. **c, e**, Coronal section of the SC showing mCherry (mC) expression (magenta) along with Chat, Cartpt, or *Otr* expression (green). AAV8 *hSyn-DIO-mCherry* (**c**) or AAV8 *hSyn-DIO-Gq-mCherry* (**e**) was used for labeling. The right graph displays quantification for efficiency (mC+/markers+) and specificity (marker+/mC+). N = 6–9 mice **d**, mC+ axons (magenta) of *Chat*+, *Cartpt*+, or *Otr*+ SPNs to DBH+ (green) areas in the CG/SMG, AM, and ST. The right graph shows the percentage of mC+ axons in the DBH+ regions. N = 4–7 mice. **f**, Coronal section of the SC showing c-Fos (green) and mC (magenta) at 2 hours after CNO administration. The right graph shows the percentage of c-Fos+ cells among mC+ cells in CNO-treated *Gq* and *mC* control mice. N = 4–10 mice. **g**, c-Fos expression (green) in the whole CG/SMG (top) and AM sections (bottom). Magenta denotes mC+ axons. The right graph shows the number of c-Fos+ cells (CG/SMG) and the percentage of c- Fos+ per DAPI+ cells (AM) in CNO-treated *Gq* and *mC* control mice. N = 7–11 mice. Data are presented as mean ± standard deviation. Statistical comparisons were conducted using one-way analysis of variance (ANOVA) followed by a two-sided Welch’s *t*-test with Bonferroni correction (**d**) or a two-sided Welch’s *t*-test (**f**, **g**). \**p* < 0.05, \*\**p* < 0.01, and \*\*\**p* < 0.001. Insets in panels **c**–**g** show magnified and channel-separated images in the boxed area. Scale bars, 100 μm for low-magnification images (**c–f**), 200 μm for low-magnification images (**g**), and 50 μm for magnified images. For more data, see Extended Data Figs. 2–4.

We next explore the functional aspects of the target specificity by injecting an AAV expressing Cre-dependent *Gq-mCherry*^27^ into the T8–T12 of *Chat-Cre* or *Cartpt-Cre* mice (Fig. 2a, b). Within the SC, *Gq-mCherry* expression was specifically observed in the marker+ SPNs (Fig. 2e). Histochemical analyses of c-Fos, an immediate early gene product serving as a proxy for cellular activation, revealed that the majority of mCherry + cells turned c-Fos+ following the administration of clozapine-N-oxide (CNO), confirming the system’s capability for specific activation of molecularly defined SPN types (Fig. 2f). Activation of *Chat*+ SPNs led to increased c-Fos expression in both the CG/SMG and AM (Fig. 2g). By contrast, activation of *Cartpt*+ SPNs induced c-Fos selectively in the CG/SMG but not in the AM (Fig. 2g). Of note, the timing of this c-Fos analysis relative to the CNO administration was determined through serial time course analysis (Extended Data Fig. 3a). Negative controls, omitting *Gq* expression, exhibited no c-Fos induction (Fig. 2f, g, right graphs, and Extended Data Fig. 3b). Collectively, these findings demonstrate that *Cartpt*+ SPNs selectively innervate the CG/SMG without affecting AM or ST, supporting a labeled line organization (Fig. 1a_1_).

## *Otr* delineates a subset of *nNos*+ SPNs projecting to the AM

Next, we sought SPNs targeting the AM. Based upon literature suggesting an association between the neural hormone oxytocin (OT) and AM functions^28–30^, along with the documented expression of *Otr* mRNA in the IML of the SC^31^, we identified *Otr*+ SPNs as specific projectors to the AM. First, utilizing the RNAscope single-molecule *in situ* hybridization method, we found that the majority of *Otr*+ cells were nNos+ in the IML of the thoracic SC (Extended Data Fig. 4a, b), consistent with the reported high-level expression of *nNos* in SPNs projecting to the AM^10^. Minimal overlap was observed between *Otr*+ and Cartpt+ cells (Extended Data Fig. 4a, b), distinguishing *Otr*+ SPNs from those projecting to the CG/SMG. We then repeated anterograde axon tracing and c-Fos induction assays shown in Fig. 2 with *Otr-Cre* mice^32^ to selectively target *Otr*+ SPNs in the T8–T12 of SC. Histochemical analyses confirmed the target specificity (Fig. 2c), and SPNs targeted by *Cartpt*-Cre and *Otr-Cre* were mutually exclusive (Extended Data Fig. 3c). In stark contrast to *Cartpt*+ SPNs, mCherry+ axons of *Otr*+ SPNs were observed in the AM, excluding the CG/SMG and ST (Fig. 2d). Similarly, when CNO was administrated into animals expressing Gq-mCherry selectively in the *Otr*+ SPNs (Fig. 2e), the majority of mCherry+ cells in the SC turned c-Fos+ (Fig. 2f). Activation of *Otr*+ SPNs led to increased c-Fos expression in the AM but not in the CG/SMG (Fig. 2g). These findings demonstrate that *Otr*+ SPNs selectively innervate the AM.

To characterize the TC type of *Otr*+ SPNs, we extracted representative marker genes for each one or few *nNos*^high^ SPN clusters (Extended Data Fig. 4c). Despite its low expression, snRNAseq data suggested the predominant presence of *Otr* in cluster 4 within the *nNos*^high^ SPNs, which was characterized by the expression of the *Palladin* (*Palld*) gene encoding a cytoskeletal-associated protein as a marker (Extended Data Fig. 4c, d). We then detected *Otr* and *Palld* expression in the IML of the T8–T13 segments and found that the vast majority of *Otr*+ cells also expressed *Palld*, and vice versa (Extended Data Fig. 4e, f). Taken together, *Otr+* SPNs provide a second example of the labeled-line organization (Fig. 1a_1_): a specific TC type of SPN projects specific output.

## Activation of *Cartpt*+ SPNs decreases intestinal motility

The presence of molecularly defined output specificity of SPNs allows the selective manipulation of sympathetic outflows. As a proof-of-concept, we next investigated whether the chemogenetic activation of *Cartpt*+ SPNs (Fig. 2e–g) influences the gastrointestinal (GI) tract, one of the major targets of postGNs in the CG/SMG^33^ (Fig. 3a, b). Red dye (carmine red) was orally administered, and the latency to the excretion of red-colored stools was measured as the GI transition time (GITT). The activation of *Chat*+ SPNs significantly increased GITT, consistent with the known negative influence of sympathetic output on the motility of the GI tract^33^. Notably, activation of *Cartpt*+ SPNs led to an increase in GITT, whereas *Otr*+ SPNs did not affect GITT (Fig. 3c). We also analyzed GITT under celiac ganglionectomy conditions (Fig. 3d). Despite the activation of *Cartpt*+ SPNs, we observed no increase in GITT in this condition, confirming that the increase in GITT induced by the activation of *Cartpt*+ SPNs is mediated through the CG/SMG (Fig. 3e). As feeding and gut contents can impact GITT^34^, we also confirmed that the activation of *Cartpt*+ SPNs increased GITT even without feeding (Fig. 3f).

**Fig. 3:**
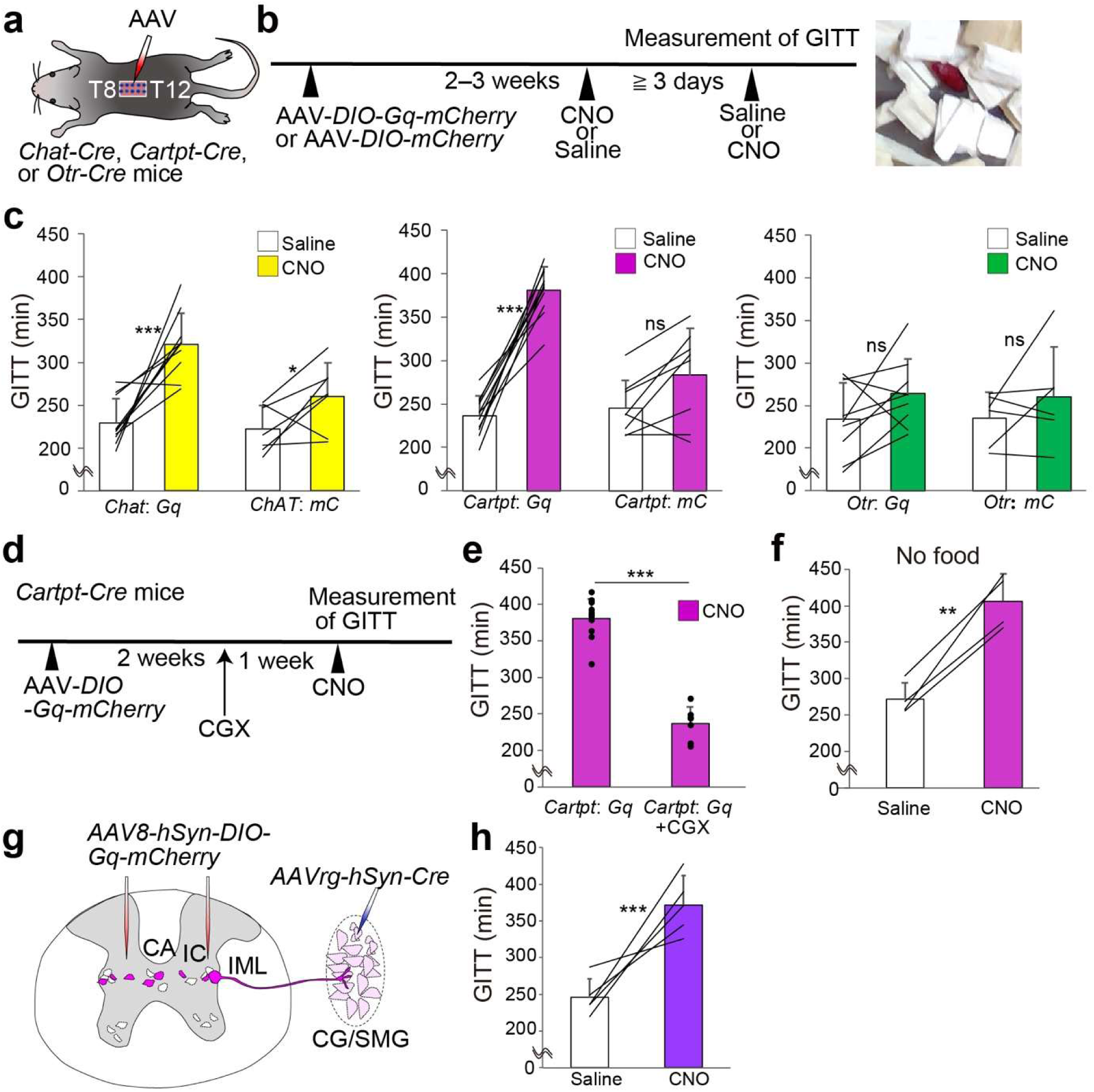
Activation of *Cartpt*+ SPNs reduces gastrointestinal motility. **a**, **b**, Schematics illustrating the experimental design (**a**) and time line (**b**). GITT, gastrointestinal transit time. The right image displays a fecal pellet stained red through carmine red administration. **c**, GITT in *Chat-Cre*, *Cartpt-Cre*, or *Otr-Cre* mice injected with AAV8-*hSyn-DIO-Gq-mCherry* or AAV8-*hSyn-DIO-mCherry* (for control), upon administration of saline (white bar) or CNO (colored bar). The results from the same mice are connected with a line in the graph. N = 6–10 mice. **d**, Schematics showing the Gq-mediated activation of *Cartpt+* SPNs under the celiac ganglionectomy (CGX) condition. **e**, GITT in CNO-treated *Cartpt-Cre* mice injected with AAV8-*hSyn-DIO-Gq-mCherry* under intact (left, N = 11) and CGX (right, N = 7) conditions. Data of intact conditions are the same as panel **c**. **f**, GITT measured without food in *Cartpt-Cre* mice injected with AAV8-*hSyn-DIO-Gq-mCherry*. N = 4 mice. **g**, Schematic showing selective activation of SPNs projecting to the CG/SMG by injecting AAV-*hSyn-DIO-Gq-mCherry* into the T8–12 SC, to which Cre is retrogradely introduced from the CG/SMG via AAVrg-*hSyn-Cre* in wild-type mice. **h**, GITT measured based on the panel **e** scheme upon administration of saline (white) or CNO (purple). N = 5 mice. Data are presented as mean ± standard deviation. Statistical comparisons were made using two-way ANOVA with repeated measures; (*Chat-Cre*) solution effect, p < 0.01, AAV effect, p < 0.01, interaction effect, ns. \**p* < 0.05, \*\*\**p* < 0.001 by a post hoc two-sided Welch’s *t*-test with Bonferroni correction. (*Carpt-Cre*) solution effect, p < 0.01, AAV effect, p < 0.01, interaction effect, p < 0.01. \*\*\**p* < 0.001 by a post hoc two-sided Welch’s *t*-test. (*Otr-Cre*) interaction effect, ns (**c**). \*\**p* < 0.01, \*\*\**p* < 0.001 by two-sided Welch’s *t*-test (**e**, **f**, **h**). For more data, see Extended Data Fig. 5.

In the thoracic SC, *Cartpt* is expressed by not only SPNs but also cholinergic interneurons^12,35^ (Extended Data Fig. 5a–d). To exclude the possibility that the observed increase in GITT by the activation of *Cartpt*+ spinal cells is caused by cholinergic interneurons, we restricted targeted *Gq-mCherry* to spinal neurons retrogradely transduced with Cre from the axons in the CG/SMG (Fig. 3g). We confirmed that exclusive activation of SPNs projecting to the CG/SMG significantly increased GITT (Fig. 3h). Collectively, these data demonstrate that activation of *Cartpt*+ SPNs predominantly suppresses the motility of the GI tract via postGNs in the CG/SMG, indicating that a specific SPN type corresponds to the regulation of the GI tract, although we do not exclude the possibility that *Cartpt*+ SPNs influence other organs (see limitations of our study in Discussion).

## *Otr*+ SPNs regulate glucose metabolism

We next sought to investigate the function of *Otr*+ SPNs. A prompt surge in blood glucose levels (BGLs) is a characteristic fight-or-flight response, providing energy supply to address impending danger or stressors^18^. Therefore, we examined BGLs upon the activation of specific SPNs. The chemogenetic activation of *Chat*+ SPNs using Gq-mCherry (Fig. 2e–g) led to a significant increase in BGLs in male mice, while such an effect was not observed in female mice (Fig. 4a, b). Although the precise reasons for this pronounced sex-specific response remain unknown, we observed that the administration of CNO induced comparable levels of c-Fos expression in the AM of both sexes (Extended Data Fig. 3d), suggesting that sex differences in BGLs occur downstream of the activation of chromaffin cells in the AM. This observation aligns with previous findings supporting sex differences in the sensitivity to sympathetic activation^36^. Notably, a similar male-specific increase in BGLs was observed upon activating *Otr*+ SPNs, but not *Cartpt*+ SPNs (Fig. 4a, b). Negative control experiments expressing *mCherry* alone had no impact on BGLs (Extended Data Fig. 6a, b). To assess whether *Otr*+ SPNs influence glucose metabolism through the AM, we examined their activation in male mice under adrenalectomy conditions (Fig. 4c). No increase in BGLs was observed following CNO administration, confirming that the rise in BGLs induced by the activation of *Otr*+ SPNs is mediated through the adrenal gland (Fig. 4d, e). These data demonstrate that the activation of *Otr*+ SPNs induces a sex-dependent increase in BGLs through the function of the AM.

**Fig. 4:**
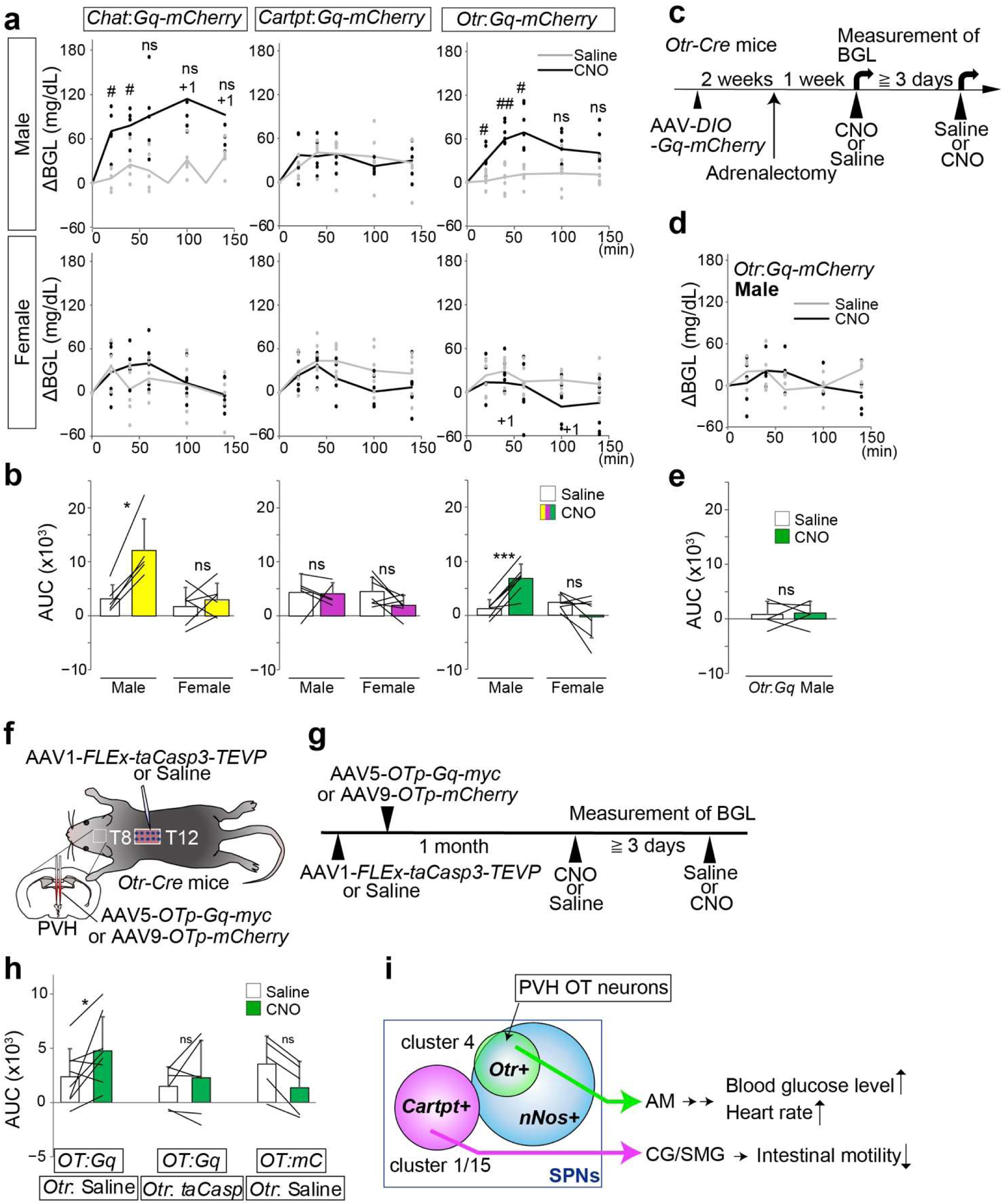
*Otr*+ SPNs mediate hypothalamic oxytocin-induced glucose metabolism. **a**, Change in blood glucose levels (ΔBGL) in *Chat-Cre*, *Cartpt-Cre*, or *Otr-Cre* mice injected with AAV8-*hSyn-DIO-Gq-mCherry* into the T8–T12 spinal segments, upon administration of saline (gray) or CNO (black). Individual data points are depicted with dots, and the average is denoted by a line. The upper graphs represent results for males, whereas the lower graphs represent results for females. +1 denotes that one data point is an outlier. **b**, Area under the curve (AUC) measurements for 0 to 140 minutes. N = 5–7 mice per condition. **c**, Schematics illustrating the Gq-mediated activation of *Otr*+ SPNs in male mice under adrenalectomy. **d**, ΔBGL under adrenalectomy, upon administration of saline (gray) or CNO (black). **e**, AUC measurements for 0 to 140 minutes. N = 5 male mice. **f**, **g**, Schematics illustrating the experimental design (**f**) and time line (**g**) for the ablation of *Otr*+ SPNs and the activation of oxytocin (OT) neurons in the paraventricular hypothalamus (PVH). *OTp*, *oxytocin* promoter. **h**, AUC measurements for 0 to 140 minutes following CNO administration. N = 5–8 mice. Data are presented as mean ± standard deviation. **i**, Schematic summary of the present work, showing that molecularly defined distinct TC types of SPNs regulate different downstream targets. Statistical comparisons were made using two-way ANOVA. Only the *Chat*: *Gq*, male and *Otr: Gq*, male groups exhibited significance; solution effect, *p* < 0.05; time course effect, *p* < 0.01; interaction effect, *p* < 0.05. #, *p* < 0.05, ##, *p* < 0.01 by a post hoc two-sided Welch’s *t*-test with Bonferroni correction showing the significant difference between the saline and CNO groups (**a**). Two-way ANOVA with repeated measures; (*Chat-Cre*) sex effect, *p* < 0.05, solution effect, p < 0.01, interaction effect, p < 0.01. \**p* < 0.05 by a post hoc two-sided Welch’s *t*-test. (*Carpt-Cre*) interaction effect, ns. (*Otr-Cre*) sex effect, *p* < 0.05, solution effect, ns, interaction effect, p < 0.01. \*\*\**p* < 0.001 by a post hoc two-sided Welch’s *t*-test (**b**). Two-sided Welch’s *t*-test (**e**), and two-way ANOVA with repeated measures; solution effect, ns, AAV effect, ns, interaction effect, *p* < 0.05. \**p* < 0.05 by a post hoc Wilcoxon’s signed-rank test (**h**). For more data, see Extended Data Figs. 6 and 7.

To ascertain whether the chemogenetic activation of *Otr*+ SPNs influences endocrinological cardiac modulations, we monitored heart rates (HRs) using a telemetry system (Extended Data Fig. 6c–g). Given that the SPNs regulating cardiac functions are located in the upper thoracic segments^37^, the Gq-mediated activation of *Otr*+ SPNs in the T8–T12 segments in this experiment should not impact them directly. While the HR in saline-injected control animals remained unaffected, a significant increase in HR was observed following CNO administration (Extended Data Fig. 6d, e). Negative control experiments expressing *mCherry* alone had no discernible impact on HR (Extended Data Fig. 6f, g). Collectively, these data suggest that manipulating the activity of *Otr*+ SPNs in the lower thoracic SC can influence metabolic and cardiac functions.

Finally, we examined the potential relationship between oxytocin (OT) produced in the hypothalamus and *Otr*+ SPNs in the lower thoracic SC. OT is a nonapeptide hormone produced by OT neurons in the paraventricular hypothalamus (PVH) that regulates energy metabolism^38^, alongside reproductive and social functions^39^. Mice lacking *OT* or the *Otr* gene exhibit obesity linked to reduced energy expenditure^40,41^. PVH OT neurons are activated during hypoglycemia, send axonal projections to the SC, and influence glucose tolerance^42^. Based on these preceding studies, we hypothesized that PVH OT neurons may contribute to the regulation of BGLs via the function of *Otr*+ SPNs in the lower thoracic SC. To test this hypothesis, we aimed to activate chemogenetically PVH OT neurons while viral-genetically ablating *Otr*+ SPNs. We introduced *OT promoter* (*OTp*) driving Gq-myc^43^, or mCherry as a control, specifically to PVH OT neurons in *Otr-Cre* male mice that received an AAV injection expressing Cre-dependent active caspase (taCasp3-TEV)^44^, or saline as a control, into the bilateral T8–T12 segments (Fig. 4f, g). Over the subsequent 1-month post-viral injection, we observed Myc-tag expression, an inference of Gq expression, in the PVH OT neurons (Extended Data Fig. 7a). Activation of PVH OT neurons increased the c-Fos expression in IML of T8–T12 segments (Extended Data Fig. 7b, c). In the mCherry control group, we also observed direct axonal projections of PVH OT neurons at the proximity of *Otr*+ SPN cell bodies in the IML, confirming the direct projection (Extended Data Fig. 7d). In the taCasp3-injected group, the number of *Otr*+ SPNs detected by RNAscope in the T8–T12 segments was significantly reduced by one-third (Extended Data Fig. 7e, f). In the saline-injected control group, where the *Otr*+ SPNs remained intact, a significant increase in BGLs was observed after the chemogenetic activation of PVH OT neurons (Fig. 4h, Extended Data Fig. 7g). By contrast, in animals with ablated *Otr*+ SPNs in the T8–T12 segments, no elevation in BGLs occurred after CNO administration. In the negative control group expressing only *mCherry* in PVH OT neurons, CNO administration did not impact BGLs. These data support the notion that the positive impact of OT on peripheral glucose metabolism is indeed mediated by *Otr*+ SPNs in the lower thoracic SC.

## Discussion

The present study established a viral-genetic approach to studying mouse SPNs in terms of their anatomical and functional organizations. The findings demonstrate the existence of molecularly delineated parallel sympathetic outflows that regulate the CG/SMG and AM (Figure 4i). Here, we discuss the biological insights provided by this study and its limitations.

While early twentieth-century physiologists accentuated the homogeneous nature of sympathetic outflows^45^, a large body of research subsequently supported the notion of selective yet coordinated control within this system^1,4,5^. For instance, classical electrophysiological studies have suggested that reflex patterns exhibited by SPNs in response to stimulation from various afferents possess unique characteristics corresponding to distinct functional pathways^5,8,9^. Under experimental conditions, animals have demonstrated the ability to favor the activation of certain selective sympathetic outflows^6,7^. Sympathetic output specificity has also been suggested in recent research regarding the somatotopic nature of acupuncture^46^, the selective sympathetic impacts on the GI tract mediated by vagal sensing of gut microbiota^33^ or gut mechanosensation^47^, and the top-down regulation of immune responses within delimited regions of the body^48^. However, the precise cellular underpinnings within SPNs for these distinctive sympathetic outflows remain largely unknown.

While classical studies have suggested the nonuniform expression of various neurochemical markers among SPNs in various species^1,6,10,11,49^, our data enhance the resolution of molecular and anatomical profiles, thereby establishing a clearer association between molecularly defined SPN types and their output specificity through three key advancements. First, we utilized recently published snRNAseq data of cholinergic SPN populations in mice^12^. This facilitated the identification of molecular markers that more accurately characterize the TC types of SPNs (Figs. 1 and Extended Data Fig. 3). Second, through a viral-genetic approach, we achieved anterograde axon mapping of genetically defined SPN types exclusively from a target SC area (Fig. 2). This not only supplements information obtained via classical retrograde approaches, but also resolves ambiguities associated with the cytotoxicity of trans-neural tracer viruses^50^, the specificity of antibodies, and difficulties in injecting tracers into a specific peripheral organ. Third, we employed a chemogenetic approach^27^ to manipulate specific SPN types directly for functional assays, a capability unavailable in classical studies. Collectively, the present study opens a new avenue for functionally classifying mouse SPNs. We believe that our approach, combining transcriptome data, mouse genetics, and viral-genetic tools, can have broad applicability to other SPN types along the entire SC.

Given that our identified markers (*Cartpt* and *Otr*) of specific SPNs represent a neural peptide and a receptor, it is intriguing to ask whether these molecular markers hold functional relevance. Despite the well-established functions of CART (encoded by *Cartpt*) in suppressing appetite in the arcuate nucleus of the hypothalamus^51,52^, the role of this peptide in the SC or the receptor mechanisms in the downstream CG/SMG remains unknown and thus constitutes an important focus for future studies. In particular, it is interesting to investigate the potential involvement of *Cartpt*+ SPNs and CART peptides in regulating the GI tract during constipation induced by the depletion of gut microbiota^33^ and diarrhea induced by the loss-of-mechanosensation of gut contents^47^. Our data, illustrating that *Otr*+ SPNs underlie OT-induced increase in BGLs (Fig. 4) lend support to the idea that OTR directly senses OT released from the PVH OT neurons and activates *Otr*+ SPNs. Future studies utilizing conditional knockout of the *Otr* gene^53^ could address this model. Given the broad involvement of OT in regulating energy expenditure^30,38,54^, an investigation into whether manipulating *Otr*+ SPNs could impact the progression of obesity represents a crucial step in establishing novel therapeutic targets for metabolic syndrome. In a broader context, selectively manipulating organ functions via specific SPN types may lay the basis for future therapeutic strategies addressing organ dysfunction associated with severe infectious disorders, traumatic SC lesions, and aging.

Finally, we acknowledge and discuss two major limitations of our study. First, we did not examine the cellular logic governing postGNs in the CG/SMG, which regulate various visceral organs in addition to the GI tract. A recent seminal study illustrated that *Cartpt*+ viscerofugal neurons in the small intestine project to the CG/SMG, and the chemogenetic activation of these neurons elevates BGLs, presumably via postGNs targeting the liver and pancreas, without affecting GITT^55^. This result sharply contrasts with our findings, where chemogenetic activation of *Cartpt*+ SPNs resulted in a specific reduction in GITT (Fig. 3) without affecting BGLs (Fig. 4). An intriguing interpretation of these results involves labeled-line connections from the *Cartpt*+ intestinal neurons to the liver/pancreas and *Cartpt*+ SPNs to the GI tract via specific postGNs in the CG/SMG, although we do not intend to exclude the possibility that *Cartpt*+ SPNs influence multiple organs. Future studies utilizing transcriptomic profiling of postGNs in the CG/SMG, similar to the approach demonstrated in the stellate ganglion^56^, along with specific anatomical/functional assays, could elucidate this proposed model. Second, our study did not examine the upstream neural circuits organized for specific types of SPNs, both within the SC and emanating from the higher brain regions^57^. In the case of skeletal motor neurons, the application of viral-genetic mapping and manipulation techniques^14^ has unveiled intricate local and long-range circuits regulating antagonistic muscles across the SC^58,59^. A similar approach could shed light on how multiple organ functions are selectively yet coordinately regulated.

## Materials and Methods

### Animals

Animals were maintained at the animal facility of the RIKEN Center for Biosystems Dynamics Research (BDR) under a 12-hour light/12-hour dark cycle with *ad libitum* access to food and water unless otherwise mentioned. Wild-type C57BL/6N mice were purchased from Japan SLC (Shizuoka, Japan). *Ai9* (known as B6.Cg-*Gt(ROSA)26Sor^tm9(CAG-tdTomato)Hze^*/J, Jax #007909) was originally purchased from the Jackson Laboratory by T. Imai, *Chat-iresCre* (also known as B6;129S6-*Chat^tm2(cre)Lowl/^*J, Jax #006410) was a kind gift from B. Lowell, and *Otr-Cre* (B6;Cg-Oxtr<em2(1xPA/T2A/icre)Yinn>, RIKEN BRC11687, also known as B6; C3-*Oxtr^em2(icre)Yinn^*/J, Jax #037578) mice were provided by Y.U. Inoue. *DBH-CreER^T2^* mice and *Cartpt-Cre* mice were originally generated in this study (as described in **Generation of knock-in mice**). All animal experiments were approved by the Institutional Animal Care and Use Committee of the RIKEN Kobe Branch.

We pooled data obtained from both male and female mice for retrograde tracing (Fig. 1), histochemical analyses (Figs. 1), axonal projections (Fig. 2), quantitative analysis of c-Fos (Fig. 2), and measurement of intestinal motility (Fig. 3). However, data on BGLs (Fig. 4) were presented separately because of the identification of a significant sex-dependent difference. Furthermore, experiments involving the measurement of HR (Extended Data Fig. 6) and the ablation of *Otr*+ SPNs (Fig. 4 and Extended Data Fig. 7) were exclusively conducted using male mice.

### Generation of knock-in mice

*DBH-CreER^T2^* (accession No. CDB0138E) and *Cartpt-Cre* (accession No. CDB0267E) lines (listed in https://large.riken.jp/distribution/mutant-list.html) were generated by CRISPR/Cas9-mediated knock-in in zygotes, as previously described^26^. For *DBH-CreER^T2^*, the *ires-CreER^T2^* cassette (*CreER^T2^* was derived from plasmid #46388, Addgene) was inserted just after the coding end of exon 12 using the homologous recombination-based method, and a mixture of crRNA (CRISPR RNA) 1 (5’-AGA AUA GCU UCU CAC AAG GUg uuu uag agc uau gcu guu uug) (50 ng/μl), tracrRNA (trans-activating crRNA) (100 ng/μl), donor vector (10 ng/μl), and Cas9 protein (100 ng/μl) was used for microinjection. For *Cartpt-Cre*, the *ires2-Cre* cassette (*ires2* was derived from plasmid #159600, Addgene) was inserted just after the coding end of exon 3 using the microhomology-mediated end joining (MMEJ)- based method, and a mixture of crRNA 2 (AGA GGG AAU AUG GGA ACC Gag uuu uag agc uau gcu guu uug) (50 ng/μl), crRNA 3 (5’-GCA UCG UAC GCG UAC GUG UUg uuu uag agc uau gcu guu uug-3’) (50 ng/μl), tracrRNA (trans-activating crRNA) (200 ng/μl), donor vector (10 ng/μl), and Cas9 protein (100 ng/μl) was used. The guide RNA (gRNA) sites were designed by using CRISPRdirect^60^ to target downstream of the stop codon. crRNA and tracrRNA (5’-AAA CAG CAU AGC AAG UUA AAA UAA GGC UAG UCC GUU AUC AAC UUG AAA AAG UGG CAC CGA GUC GGU GCU) were purchased from FASMAC (Atsugi, Japan).

For *DBH-CreER^T2^* mice, 38 F_0_ founder mice were obtained from 230 transferred zygotes, 32 of which were *Cre*-positive as identified by PCR. The line was established by one male, which harbors a targeted sequence identified by PCR and sequencing. PCR was performed using the following primers, as shown in Fig. 1:

*- DBH-WT-F* 5’-GGAAGAGGTGTGGAGTGTTCAGCATGGGAG
*- ERT2-F* 5’-CACTGCGGGCTCTACTTCATCGCATTCC
*- DBH-WT-R* 5’-GAGCTGTCCCTATGTGGAGTCAGCAGTGTG

For the sequence analysis, the PCR products from using the primer *DBH-WT-F* and *DBH-3’junc-R* 5’- TCTGAAGAATAGCTTCTCACAAgagctcag were subcloned into the *pCR Blunt II TOPO* vector (Zero Blunt TOPO PCR Cloning Kit, Thermo Fisher Scientific) and sequenced using *M13-Foward* and *M13-Reverse* primers.

For *Cartpt-Cre* mice, 37 F_0_ founder mice were obtained from 218 transferred zygotes, 13 of which were *Cre*-positive as identified by PCR and sequencing. The line was established by one male, which harbors a targeted sequence identified by PCR and sequencing. PCR was performed using the following primers, as shown in Extended Data Fig. 3:

*- Cartpt-F* 5’-CGGATCTGACTGCTTCGACCTGAGC
*- Cre-F* 5’-GTCGAGCGATGGATTTCCGTCTCTGG
*- Cartpt-R* 5’-GAAGCAACAGGGAAAGAGCCCATCCG

For the sequence analysis, the PCR products from using the primer *Cartpt-F* and *Cartpt-3’ junk-R* 5’- GTAAGCTGAGGTGAAGCCAGACATgtcgac or *Cartpt-5’ junk-F* 5’- AGAGGGAATATGGGAACgaattcgc and *Cartpt-R* were subcloned into the *pCR Blunt II TOPO* vector (Zero Blunt TOPO PCR Cloning Kit, Thermo Fisher Scientific) and sequenced using *M13-Foward* and *M13-Reverse* primers.

The germline transmission of these mouse lines was confirmed by genotyping of the F_1_ mice. Of note, consistent expression patterns with the endogenous genes were confirmed by the expression of the Cre reporter fluorescent protein for *DBH-CreER^T2^*(Extended Data Fig. 1) and the histochemical analysis for *Cartpt-Cre* (Fig. 2). For *DBH-CreER^T2^*mice, tamoxifen (Sigma, #10540-29-1) was dissolved in corn oil at 20 mg/ml by shaking overnight at 37 °C and stored at 4 °C for injections. A dosage of 75 mg/kg of tamoxifen (from a 20 mg/ml solution) was administered intraperitoneally to induce the activation of CreER^T2^.

### Virus

The following AAV vectors were obtained from Addgene:

*-* AAV2retro *hSyn.Cre.WPRE.hGH* (1.8 × 10^13^ gp/ml) (Addgene #105553)
*-* AAV8 *hSyn-DIO-hM3D(Gq)-mCherry* (2.1 × 10^13^ gp/ml) (Addgene #44361)
*-* AAV8 *hSyn-DIO -mCherry* (2.3 × 10^13^ gp/ml) (Addgene #50459)

The following AAV vector was obtained from the UNC viral core:

*-* AAV1 *EF1a-FLEx-taCasp3-TEVp* (5.8 × 10^12^ gp/ml)

The following AAV vector was constructed by K. Inada^43^ and generated by the UNC vector core:

*-* AAV5 *OTp-hM3Dq-Myc* (2.4 × 10^13^ gp/ml)

The following AAV vector was constructed by K. Inada^43^ and generated by the Viral Vector Core, Gunma University Initiative for Advanced Research:

*-* AAV9 *OTp-mCherry* (2.9 × 10^13^ gp/ml)

### Retrograde tracing from the CG/SMG

*Ai9* mice (aged 3–4 weeks) underwent anesthesia through the intraperitoneal injection of a combination of 65 mg/kg ketamine (Daiichi-Sankyo) and 13 mg/kg xylazine (Sigma-Aldrich). For the CG/SMG (Fig. 1), an incision was made along the midline, and saline-soaked gauze was placed on the abdomen. To expose the CG/SMG, adjacent organs such as the stomach and intestines were carefully relocated to the moist gauze on the side of the mouse. Consistent moistening with saline was maintained during the surgical procedure to prevent dehydration. The CG/SMG, positioned around the descending aorta and left renal artery, surrounding the superior mesenteric artery, was identified.

A glass pipette prepared for a microinjector (#UMP3, WPI) was utilized to administer 500 or 100 nl of AAVrg *hSyn-Cre-WPRE-hGH* (Addgene #105553) with 0.1% Fast Green (#F7252, Sigma-Aldrich) into the CG/SMG of *Ai9* mice, respectively, at an infusion rate of 50 nl per minute^33^. Successful injection was confirmed by observing the ganglia staining with Fast Green. Following the injection, the abdominal cavity was rinsed with saline to prevent AAV dispersion, and saline introduction into the abdominal cavity post-surgery aimed to prevent adhesion of the abdominal organs. The incisions were then sutured. Following the surgery, an antipyretic analgesic, ketoprofen (5 mg/kg, Kissei Pharmaceutical) was administered intramuscularly, and an antibiotic, gentamicin (5 mg/kg, Takada Pharmaceutical), was administered intraperitoneally. Approximately 2 weeks after the surgical procedure, the mice were anesthetized with isoflurane, euthanized, and subsequently perfused with phosphate-buffered saline (PBS), followed by a 4% paraformaldehyde (PFA) solution in PBS. The SC was post-fixed with a 4% PFA solution in PBS.

For the tissue clearing of the CG/SMG (Supplementary Movie 1) and SC (Fig. 1g and Supplementary Movie 2), the CUBIC method^21^ was implemented. Specifically, after perfused fixation, the specimen underwent overnight fixation in a 4% PFA solution in PBS. Subsequently, the specimen was washed with PBS for at least 2 hours, repeated twice, at 25 °C. The fixed tissues were then immersed in 5 ml of half-diluted CUBIC-L, which is a mixture of 10% (wt/wt) N-butyl diethanolamine and 10% (wt/wt) Triton X-100 in deionized water, with gentle shaking for at least 6 hours at 37 °C. The solution was then replaced with 5 ml of fresh CUBIC-L, and gentle shaking at 37 °C was continued for an additional 1–2 days. To terminate the delipidation reaction, the samples were washed with 5 ml PBS, again with gentle shaking, at 25 °C for at least 2 hours. This washing step was repeated more than three times. Subsequently, the samples were immersed in 5 ml of half-diluted CUBIC-R+, which is a mixture of 45% (wt/wt) antipyrine, 30% (wt/wt) nicotinamide, and 0.5% (vol/vol) N-butyl diethanolamine in deionized water, with gentle shaking at 25 °C for at least 6 hours. The sample was further immersed in 5 ml of CUBIC-R+ placed in a 5 ml tube and gently shaken at 25 °C overnight.

The cleared CG/SMG or SC samples were carefully placed into glass capillaries and then positioned in the Zeiss Lightsheet Z.1 Light Sheet Fluorescence Microscope. Z-stack images were obtained using a 5× objective lens (numerical aperture [NA]: 0.1). Subsequently, the captured images were adjusted post hoc using Imaris 9.8 software, and three-dimensional reconstructions were saved as MP4 video files (Supplementary Movies 1 and 2).

For the examination of marker gene expression in retrogradely labeled cells, AAVrg *hSyn-Cre-WPRE-hGH* was administered into the CG/SMG of *Ai9* mice. At approximately 2 weeks post-injection, perfused fixation was carried out. The percentage of specific gene expression was calculated based on 25–70 tdTomato+ cells in the CA, IC, and IML of SC sections. Detained information on the procedures can be found in the **Histochemistry** section.

### Histochemistry

For immunostaining, after conducting perfused fixation of the mice using a 4% PFA solution in PBS, the tissues were subjected to an additional overnight post-fixation in 4% PFA in PBS at 4 °C. The specimens were cryoprotected with a 30% sucrose solution in PBS at 4 °C for 24 hours, and then embedded in the OCT compound (#4583; Tissue-Tek). Next, 30-μm coronal sections were obtained using a cryostat (model #CM1860; Leica) and placed on MAS-coated glass slides (Matsunami). The specimens were subjected to a triple wash with PBS and treated with a 5% normal donkey serum (NDS; Southern Biotech, #0030-01) solution in PBST (PBS with 0.1% Triton X) for 30 minutes at 25 °C for blocking. The following primary antibodies were used in this study: goat anti-mCherry (at a dilution of 1:250; #AB0040-200, SICGEN), rabbit anti-DBH (at a dilution of 1:1000; #ab209487, Abcam), goat anti-Chat (at a dilution of 1:100; #AB144P, Sigma), rabbit anti-nNos (at a dilution of 1:250; #61-7000, Thermo Fisher Scientific), goat anti-CART (at a dilution of 1:200; #AF163, R&D Systems), rabbit anti-c-Fos (at a dilution of 1:1000; #2250S, Cell Signaling), mouse anti-Myc (at a dilution of 1:500; #sc-40, Santa Cruz), and rabbit anti-OT (at a dilution of 1:500, #20068, Immunostar). Before staining using anti-DBH, antigen retrieval was conducted in Tris-Cl EDTA pH9.0. These primary antibodies were diluted in 1% NDS in PBST and applied to the sections overnight at 4 °C. These sections were then washed three times with PBS and treated with the following secondary antibodies, diluted in 1% NDS in PBST and containing 4’,6-diamidino-2-phenylindole dihydrochloride (DAPI), for 1 hour at 25 °C: donkey anti-rabbit AF488 (at a dilution of 1:200; #A32790, Thermo Fisher Scientific), donkey anti-goat AF555 (at a dilution of 1:200; #A32816, Thermo Fisher Scientific), donkey anti-mouse AF555 (at a dilution of 1:200; #A32773, Thermo Fisher Scientific), and donkey anti-rabbit AF488 (at a dilution of 1:200; #A32790, Thermo Fisher Scientific). The sections were washed three times with PBS and mounted with cover glass using Fluoromount (#K024; Diagnostic BioSystems).

*In situ* hybridization (ISH) was conducted as described previously^61^. To generate cRNA probes, DNA templates were amplified by PCR from SC cDNA (#MD-23; Genostaff). A T3 RNA polymerase recognition site (5’-AATTAACCCTCACTAAAGGG) was added to the 3’ end of the reverse primers. The primer sets and sequences of the probe targets were as follows:

*- Cartpt*-1: 5’-ggacatctactctgccgtgg; 5’-tccgggttgtgatgtcatct
*- Cartpt*-2: 5’-gccctggacatctactctgc; 5’-tccgggttgtgatgtcatct
*- Chodl*: 5’-aagggaaggcagctgcttag; 5’-aactgggagctgcttccatc
*- Pitx2*: 5’- tggaccaaccttacggaagc; 5’-aaacatttgtgggcgtacgc

DNA templates (500–1000 ng) that were amplified by PCR were then subjected to *in vitro* transcription with either DIG-(#11277073910) or Flu- (#11685619910) RNA labeling mix and T3 RNA polymerase (cat#11031163001) according to the manufacturer’s instructions (Roche). In the case of *Cartpt*, a combination of two probes was utilized to enhance the signal-to-noise ratio.

For single-color ISH combined with anti-mCherry staining, following the hybridization and washing steps, sections were incubated with a solution containing horseradish peroxidase (HRP)- conjugated anti-Dig (at a dilution of 1:500; #11207733910, Roche Applied Science) and goat anti-mCherry (at a dilution of 1:250; #AB0040-200, SICGEN) antibodies overnight. The signals were amplified by a 1:70 TSA-plus Cyanine 3 (#NEL744001KT; AKOYA Bioscience) for 25 minutes, followed by washing. mCherry-positive cells were then visualized using donkey anti-goat AF555 (at a dilution of 1:200; #32816, Invitrogen). PBS containing 50 ng/ml of DAPI (#8417, Sigma-Aldrich) was used for counter-nuclear staining.

For dual-color ISH, HRP-conjugated anti-flu antibody (at a dilution of 1:250; #NEF710001EA, AKOYA Bioscience) was used to detect Flu-labeled RNA probes by a 1:70 TSA-plus biotin (#NEL749A001KT; AKOYA Bioscience), which was then visualized with streptavidin-AF488 (at a dilution of 1:250; Life Technologies). After inactivation of HRP with a 2% sodium azide solution in PBS for 15 minutes at 25 °C, HRP-conjugated anti-Dig (at a dilution of 1:500; #11207733910, Roche Applied Science) and a 1:70 TSA-plus Cyanine 3 were used to detect Dig-labeled cRNA probes. The sections were then mounted with a cover glass using Fluoromount (#K024; Diagnostic BioSystems).

To detect the expression of *Otr* or *Palld*, we used the RNAscope Multiplex Fluorescent Reagent Kit v2 (#323100; ACD) following the manufacturer’s instructions. For single-color detection of *Otr*, we utilized specific probes for *Otr* (Mm-Oxtr #412171, ACD). To conduct dual-color imaging of *Otr* and *Palld*, we used C1-probe for *Palld* (Mm-Palld #822611, ACD) and C2-probe for *Otr* (Mm-Oxtr-C2 #412171-C2, ACD). The sections were air-dried and then incubated with DEPC-treated PBS for 5 minutes. Subsequently, they were heated in a dry oven at 60 °C for 30 minutes. The sections were fixed with a 4% PFA solution in PBS for 15 minutes at 4 °C, followed by dehydration in 100% ethanol for 5 minutes and an additional 5 minutes of air-drying. All sections were pretreated with hydrogen peroxide for 10 minutes at 25 °C, followed by washing with Milli-Q for 5–10 minutes; this step was repeated twice. Target retrieval was carried out by placing the slides in a steamer at a temperature exceeding 99 °C for 15 minutes, followed by a quick wash with Milli-Q for 15 seconds. The sections were dehydrated in 100% ethanol for 3 minutes and heated in a dry oven at 60 °C for 5 minutes. Subsequently, the sections received treatment with protease reagent (protease III) for 30 minutes at 40 °C, followed by another wash with Milli-Q. Sections were then incubated in the RNAscope probes for 2 hours at 40 °C. Following this incubation, the slides were washed with a wash buffer. The sections were then incubated in three amplification reagents at 40 °C: AMP1 and AMP2 for 30 minutes each, and AMP3 for 15 minutes. This was followed by horseradish peroxidase-C1 (HPR-C1) amplification at 40 °C for 15 minutes.

For a single-color RNAscope for *Otr* combined with anti-mCherry immunostaining, sections were subsequently incubated in TSA-plus Cyanine 3 (#NEL744001KT, AKOYA Bioscience) at a 1/1500 concentration for 30 minutes at 40 °C. In the final step of the process, sections were subjected to HRP blocking for 15 minutes at 40 °C. After the final wash, the slides were prepared for IHC to detect mCherry. Following a blocking step with 5% NDS, sections were incubated overnight with goat anti-mCherry antibody (at a dilution of 1:250; #AB0040-200, SICGEN) in 1% NDS at 4 °C. After washing three times, the sections were incubated for 1 hour with donkey anti-goat AF555 (at a dilution of 1:200; #32816, Thermo Fisher Scientific) and DAPI in 1% NDS. Subsequently, the sections were washed with PBS three times and mounted with a cover glass using Fluoromount (#K024; Diagnostic BioSystems).

For a dual-color RNAscope for *Otr* and *Palld*, combined with anti-Chat immunostaining, sections following HPR-C1 treatment were incubated with TSA-plus Cyanine 3 (#NEL744001KT; AKOYA Bioscience) at a 1/1500 concentration for 30 minutes at 40 °C, followed by HRP blocking for 15 minutes at 40 °C. After washing, sections were included with horseradish peroxidase-C2 (HPR-C2) at 40 °C for 15 minutes. Subsequently, sections were incubated in TSA-plus biotin (NEL749A001KT; AKOYA Bioscience) at a 1/1500 concentration for 30 minutes at 40 °C. Following HRP blocking for 15 minutes at 40 °C and washing, sections were subjected to a blocking step with 5% NDS and then incubated overnight at 4 °C with goat anti-Chat (at a dilution of 1:250; #AB144P, Sigma) and Streptavidin-AF488 (at a dilution of 1:250; #S32354, Thermo Fisher Scientific) in 1% NDS. After washing three times, the sections were incubated for 1 hour with donkey anti-Goat AF647 (at a dilution of 1:250; #A-21447, Thermo Fisher Scientific), and DAPI in 1% NDS. Subsequently, the sections were washed with PBS three times and mounted with a cover glass using Fluoromount (#K024; Diagnostic BioSystems).

To count the c-Fos+ cells in the whole mount preparation of CG/SMG (Fig. 2), 3 days before perfused fixation, a 300-μl solution containing 4 mg/ml of Fluorogold (#526-94003; Fluorochrome, LLC) in saline was administered via intraperitoneal injection to visualize the abdominal ganglia. The CG/SMG was dissected and subsequently fixed in a 4% PFA solution in PBS overnight. The specimen was trimmed under a fluorescence microscope using ultraviolet light to visualize Fluorogold. It was then subjected to triple washes with PBS and permeabilization using a solution comprising 0.5% Triton X-100, 0.05% Tween-20, and 4 μg/ml of heparin (referred to as PTxwH), followed by incubation overnight at 25 °C. Subsequently, primary antibodies, rabbit anti-c-Fos (at a 1:1000 dilution; #2250S, Cell Signaling) and goat anti-mCherry (at a 1:250 dilution, #AB0040-200, SICGEN), were introduced into the whole CG/SMG immersed in PTxwH and incubated at 4 °C for 3 days. Following the triple wash steps with PTxwH, the specimen was stained using donkey anti-rabbit AF488 (at a 1:200 dilution, #A32790, Invitrogen) and donkey anti-goat AF555 (at a 1:200 dilution, #A32816, Invitrogen) at 4 °C for 3 days. Triple washes with PTxwH followed this staining process. The CG/SMG specimen was then placed on a glass-bottom dish (#3960-035; IWAKI) and covered with Fluoromount (#K024; Diagnostic BioSystems).

The sections were imaged using an Olympus BX53 microscope equipped with a 10× objective lens (NA: 0.4). For confocal imaging of whole mount preparations, an inverted Zeiss LSM780 confocal microscope with a Plan-Apochromat 10× objective lens (NA: 0.45) was used. Z-stack images were acquired and subsequently processed into maximum projection images. The acquired images were processed using Fiji software. For the analysis of RNAscope data (*Otr* and *Palld*), we defined positive cells as those exhibiting five or more RNAscope dots, as the presence of a small number of dots could be observed even in sections derived from *Otr* knockout mice^53^.

### Reanalysis of snRNAseq data

The snRNAseq data sourced from spinal cord autonomic and skeletal motor neurons (accession No. GSE161621)^12^ were subjected to reanalysis in R (version 4.3.1) and RStudio (Server 2023.06.1 Build 524; supported by the RIKEN BDR DNA Analysis Facility at the Laboratory for Developmental Genome System) using Seurat (version 5.0.1)^62^. Specifically, the analysis focused on *Chat+* clusters, excluding *B cell leukemia/lymphoma 6* (*Bcl6*)+ motor neurons and (*Pax2*)+ cholinergic interneurons, narrowing the investigation to cholinergic SPNs. The standard Seurat data analysis workflow encompassed quality control, where genes were retained if at least one unique molecular identifier was detected in a minimum of three cells, followed by normalization. Subsequently, SCTransform scaling was employed, and the top 3000 variable features were identified and utilized as input for principal component analysis. The initial clustering was performed using the top 50 principal components (PCs) at a resolution of 0.05 to distinguish between neuronal and non-neuronal cell populations. Non-neuronal cells were excluded based on the expression of *Solute carrier family 7 member 10* (*Slc7a10*) and *Myelin oligodendrocyte glycoprotein* (*Mog*). The subsetUmapClust function was then applied to perform re-clustering. Furthermore, cholinergic SPNs were segregated based on the expression of *Chat*, *Bcl6*, and *Pax2* using the top 35 PCs at a resolution of 0.1. As shown in Fig. 2, SPNs were clustered using the top 50 PCs at a resolution of 0.5. We defined *nNos*^low^ clusters as those in which 50% or fewer cells expressed *nNos*, with average log-normalized expression levels (in arbitrary units) lower than zero. The remaining SPN clusters were categorized as *nNos*^high^.

### Viral injections into the SC or PVH

For viral injections into the lower thoracic SC (Figs. 2–4), *Chat-Cre*, *Cartpt-Cre*, or *Otr-Cre* mice (aged 4–5 weeks) underwent anesthesia through the intraperitoneal injection of a combination of 65 mg/kg ketamine (Daiichi-Sankyo) and 13 mg/kg xylazine (Sigma-Aldrich). An incision around the thoracic region of the SC was made. To expose the SC, the paraspinal muscles and dorsal vertebral elements were carefully excised using fine forceps. A glass pipette prepared for a microinjector (#UMP3; WPI) was utilized to administer 100 nl of the AAV vector with 0.1% Fast Green (#F7252, Sigma-Aldrich) at 10 distinct sites (infusion rate: 60–70 nl/min) bilaterally in the T8–T12 segments. In Figs. 2–4 experiments, we targeted T8–12, because SPNs in T13 and lumber region is known to project to sympathetic ganglia located lower the CG/SMG, such as inferior mesenteric ganglia^22^.

For the chemogenetic activation of PVH OT neurons (Fig. 4), *Otr-Cre* male mice (aged 6–8 weeks) underwent anesthesia as mentioned above and were secured to stereotactic equipment (Narishige). To target the AAV into the PVH, stereotactic coordinates were determined based on the Allen Mouse Brain Atlas^63^, and the following coordinates were used (measured in mm from the Bregma for anteroposterior [AP] and mediolateral [ML], and from the brain surface for dorsoventral [DV]): AP: –0.8, ML: 0.2, and DV: 4.5. The volume of AAV serotype 5 *pAAVOTp-hM3Dq-Myc* injected was 200 nl, administered at a rate of 50 nl per minute. Following the surgical procedures, an antipyretic analgesic, ketoprofen (5 mg/kg, Kissei Pharmaceutical), was administered intramuscularly, and an antibiotic, gentamicin (5 mg/kg, Takada Pharmaceutical), was administered intraperitoneally.

### Measurement of axons and c-Fos+ cells

For the measurement of axons (Fig. 2), 30-μm sections of the sympathetic ganglia and adrenal glands were prepared and immunostained with anti-DBH and anti-mCherry as described in the **Histochemistry** section. Images of these sections were acquired using an Olympus BX53 microscope equipped with a 10× objective lens (NA: 0.4). The automatic counting of mCherry-positive axons within DBH-positive cells was carried out using Fiji software. Binarized images of DBH and mCherry were generated, and the area of mCherry-positive axons was normalized to the DBH-positive area within the image. The average counts from three sections were calculated for individual *Chat-Cre*, *Cartpt-Cre*, or *Otr-Cre* mice.

To quantify the number of c-Fos+ cells in the sections of the adrenal gland and SC (Fig. 2), immunostaining was carried out as described in the **Histochemistry** section. For the adrenal gland sections, the counting of c-Fos- and DAPI-positive cells was carried out automatically using Fiji software. The noise was removed through the “Remove Outliers” function, and the count of both c-Fos- and DAPI-positive cells was determined using the “Find Maxima” function. The average number of c-Fos-positive cells across three sections was divided by the count of DAPI-positive cells in the corresponding area. This quotient was used as the value for individual mice. For the SC sections, the quantification of c-Fos-positive cells was carried out manually. Within the T8–T12 segments, the number of c-Fos-positive cells among mCherry-positive cells was counted for five sections (one section per segment,) and the counts were averaged for individual mice.

The identification and counting of c-Fos+ cells in the whole mount CG/SGM (Fig. 2) were performed automatically using Fiji software. After removing noise through the “Remove Outliers” function, the total number of Fos+ cells was determined using the “Find Maxima” function. Of note, data were not normalized to area or volume. Each data point represents the number of c-Fos+ cells per CG-SMG.

### Measurements of gastrointestinal motility and blood glucose levels

Neural activation experiments targeting hM3D (Gq) to the desired cell types (Figs. 2–4) were conducted at least 2 weeks after the AAV injection. To induce activation, a dosage of 2 mg/kg of CNO (#C0832, Sigma-Aldrich) was injected into the mice via intraperitoneal administration. Saline was injected into the same mice for the negative control. The order of CNO or Saline administration was randomized.

To measure GITT, the mice received an oral gavage of a 6% carmine red solution that was dissolved in 0.5% methylcellulose, and prepared with sterile saline. GITT was determined by measuring the time between the oral gavage and the excretion of a carmine-containing fecal pellet^33^. All mice had *ad libitum* access to food and water during the measurement period. Under the “no food condition” (Fig. 3f), food was removed just after the administration of CNO or saline. These experiments were performed during the light phase.

For celiac ganglionectomy (CGX, Fig. 3d), the CG/SMG was exposed and carefully dissected with fine forceps. Following the surgery, saline was introduced into the abdominal cavity to prevent adhesion of the abdominal organs, and the incisions were sutured. Following the surgery, ketoprofen (5 mg/kg, Kissei Pharmaceutical) was administered intramuscularly and gentamicin (5 mg/kg, Takada Pharmaceutical) intraperitoneally. At 1 week following the surgery, GITT was measured as mentioned above.

To prevent a potential stress-induced increase in BGLs, the mice underwent a 2-day habituation period before the assays. Each mouse was individually housed, and a daily 10-minute habituation was conducted to replicate the process of blood collection from the tail tip. All mice had *ad libitum* access to food and water throughout the habituation and measurement periods. To minimize the impact of stress further, the tail tip was gently incised at 2 hours before the measurement, followed by another 10-minute habituation session. Blood samples were obtained from the tail tip, and BGLs were assessed using a blood glucose meter for laboratory animals (cat# SUGL-001; ForaCare, Tokyo) at intervals of 0, 20, 40, 60, 100, and 140 minutes after the intraperitoneal administration of 2 mg/kg CNO or saline. In Fig. 4, the baseline BGL at time 0 minutes was used to calculate ΔBGL values. Of note, the baseline was not significantly different between sexes or among those injected with different AAVs (mg/dL): mean ± standard deviation, 153.70 ± 15.62 (*Chat*: *Gq,* male), 161.83 ± 19.62 (*Chat*: *Gq,* female), 143.14 ± 16.39 (*Chat*: *mC,* male), 141.83 ± 10.92 (*Chat*: *mC,* female), 143.30 ± 16.02 (*Cartpt*: *Gq,* male), 149.07 ± 24.07 (*Cartpt*: *Gq,* female), 139.50 ± 22.98 (*Cartpt*: *mC,* male), 146.88 ± 17.55 (*Cartpt*: *mC,* female), 167.43 ± 16.30 (*Otr*: *Gq,* male), 156.21 ± 25.48 (*Otr*: *Gq*, female), 162.83 ± 21.98 (*Otr*: *mC,* male), and 149.17 ± 22.24 (*Otr*: *mC*, female). In Fig. 4a, the following data points are denoted as +1 symbols in the graph as outliers: *ChAT*: *Gq*, male, 100 min; 221mg/dL, 140 min; 226 mg/dL, *Otr*: *Gq*, female, 40 min; –78 mg/dL, 100 min; –71 mg/dL.

To perform adrenalectomy (Fig. 4c–e), *Otr-Cre* male mice (aged 5–7 weeks) were anesthetized using a combination of 65 mg/kg ketamine (Daiichi-Sankyo) and 13 mg/kg xylazine (Sigma-Aldrich) via intraperitoneal injection. An incision was made along the midline, and gauze soaked in saline was placed on the abdomen. To expose the adrenal gland, adjacent organs such as the stomach and intestines were carefully relocated to the moist gauze on the side of the mouse. Throughout the surgical procedure, these organs were consistently moistened with saline to prevent dehydration. Using fine small scissors, the bilateral adrenal glands were carefully excised from the kidneys. After the surgery, ketoprofen (5 mg/kg, Kissei Pharmaceutical) was administered intramuscularly and gentamicin (5 mg/kg, Takada Pharmaceutical) intraperitoneally. Considering that the adrenal glands secrete aldosterone to maintain the salt balance, the drinking water was supplemented with a 1% w/v sodium chloride solution^64^. At 1 week following the surgery, BGLs were measured as mentioned above. The baseline was 182.10 ± 14.20 mg/dL.

### Telemetry system

To monitor the HR in mice, as shown in Extended Data Fig. 6c–g, we conducted surgical implantations of telemetry devices (G2-HR E-Mitter; Starr Life Sciences Corp., Oakmont, PA, USA) following a recovery period of at least 1 month after the AAV injection. For the implantation procedure, mice underwent intraperitoneal injection of 65 mg/kg ketamine and 13 mg/kg xylazine for anesthesia. The telemeter was placed in the abdominal cavity, with the negative lead sutured subcutaneously to the right subclavian greater pectoralis muscle and the positive lead to the chest wall on the left side of the sternum, anterior to the last rib. Following the surgery, gentamicin (5 mg/kg, Takada Pharmaceutical) was promptly administered. The mice were then maintained in their home cages for 1 week to facilitate recovery. During this recovery period, the mice were habituated to intraperitoneal injection for at least 3 days. On the recording day, saline was administered at zeitgeber Time 2 (ZT2, 10:00) and CNO at ZT6 (14:00). The signals transmitted by the telemetry device were received and recorded at 1-minute intervals using an ER-4000 receiver and VitalView software (Starr Life Sciences Corp.). The average HR value was calculated every 10 minutes. As shown in Extended Data Fig. 6d, f, the mean baseline HR during the period from –30 minutes to 0 minutes was used to calculate the ΔHR. Of note, the baseline was not significantly different between saline and CNO or those injected with different AAVs (rpm): 551.32 ± 71.70 (*Otr*: *Gq*, saline), 544.29 ± 73.05 (*Otr*: *Gq*, CNO), 565.97 ± 81.14 (*Otr*: *mC*, saline), and 594.83 ± 56.03 (*Otr*: *mC*, CNO).

### Statistical analysis

The statistical details of each experiment, including the statistical tests used, and the exact value of the number of animals are described in each figure legend. *P* values < 0.05 were considered to indicate statistical significance.

## Data and materials availability

All data are available in the main paper and supplementary materials. All materials, including the *DBH-CreER^T2^*mice and *Cartpt-Cre* mice and plasmids, are available through reasonable request to the corresponding authors.

## Code availability statement

No original code was generated in the course of this study.

## Author contributions

Y.H. and K.M. conceived the experiments. Y.H. performed the experiments and analyzed the data, with technical support by S.U., K.I., M.H., and S.I. M.T. conducted a reanalysis of the snRNAseq data. *DBH-CreER^T2^* and *Cartpt-Cre* mice were generated by M.S., T.A., and M.H., Y.U.I., and T.I. provided the *Otr-iCre* mice. Y.H. and K.M. wrote the paper with contributions from all coauthors.

## Acknowledgments

We thank the staff at the RIKEN BDR animal facility for the animal care and *in vitro* fertilization, Hideki Enomoto (Kobe University), Takeshi Imai (Kyushu University), and members of the Miyamichi Laboratory for the critical reading of the manuscript, Addgene, the University of North Carolina Vector Core, and the Viral Vector Core of Gunma University Initiative for Advanced Research for the AAV productions, and the Laboratory for Developmental Genome System for support with the snRNA-seq reanalysis. This study was supported by the RIKEN Special Postdoctoral Researchers Program and JSPS KAKENHI (21K1521) to Y.H., and the JST CREST Program (JPMJCR2021), JSPS KAKENHI (20K20589 and 21H02587), grants from Uehara Memorial Foundation, G-7 Scholarship Foundation, and RIKEN BDR QMIN project grant to K.M.

## Competing interests

The authors declare that they have no competing interests.

## Extended data figures and legends

**Extended Data Fig. 1:**
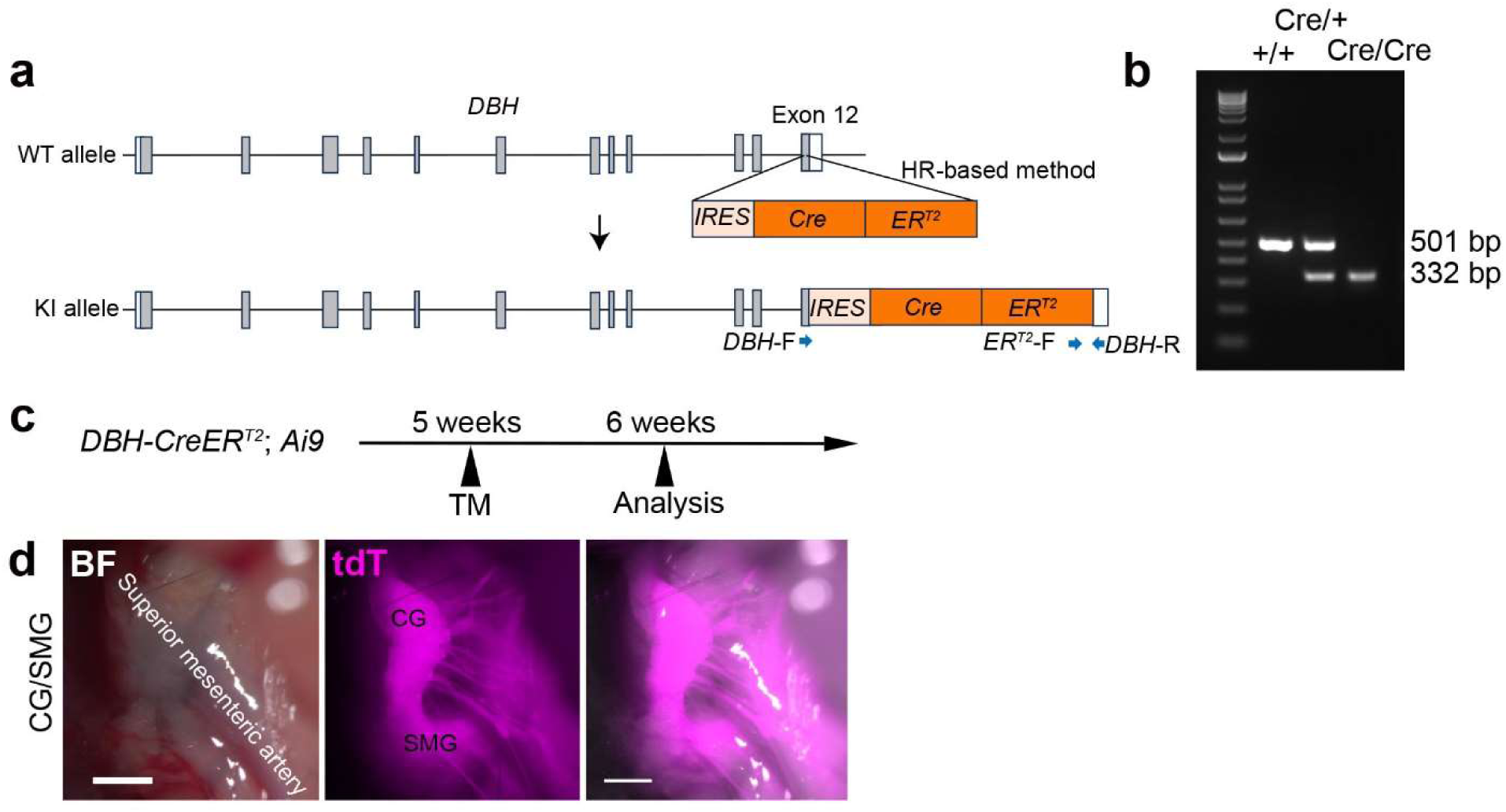
Labeling post-GNs in the CG/SMG via *DBH-CreER^T2^* mice, related to Fig. 1. **a**, Illustration of the knock-in (KI) strategy for generating *dopamine-β-hydroxylase* (*DBH*)-*CreER*^T2^ mice using the homologous recombination (HR)-based method^26^. The *DBH* gene comprises 12 exons, with open and closed boxes denoting untranslated and protein-coding regions, respectively. The KI allele is schematically depicted below, where the *IRES-Cre-ER^T2^* cassette is inserted into the coding end of the *DBH* locus located in exon 12. Blue arrows indicate the site of genotyping primer sites. **b**, Representative gel image of electrophoresis examining the *Cre* allele and a negative control (wild-type animal). **c**, Time line of the experiments for genetically labeling postganglionic neurons. TM, tamoxifen. **d**, Fluorescence microscope imaging of the CG/SMG in *DBH-CreER^T2^; Ai9* mice. BF, bright field. tdT, tdTomato from the *Ai9* allele. Scale bar, 1 mm.

**Extended Data Fig. 2:**
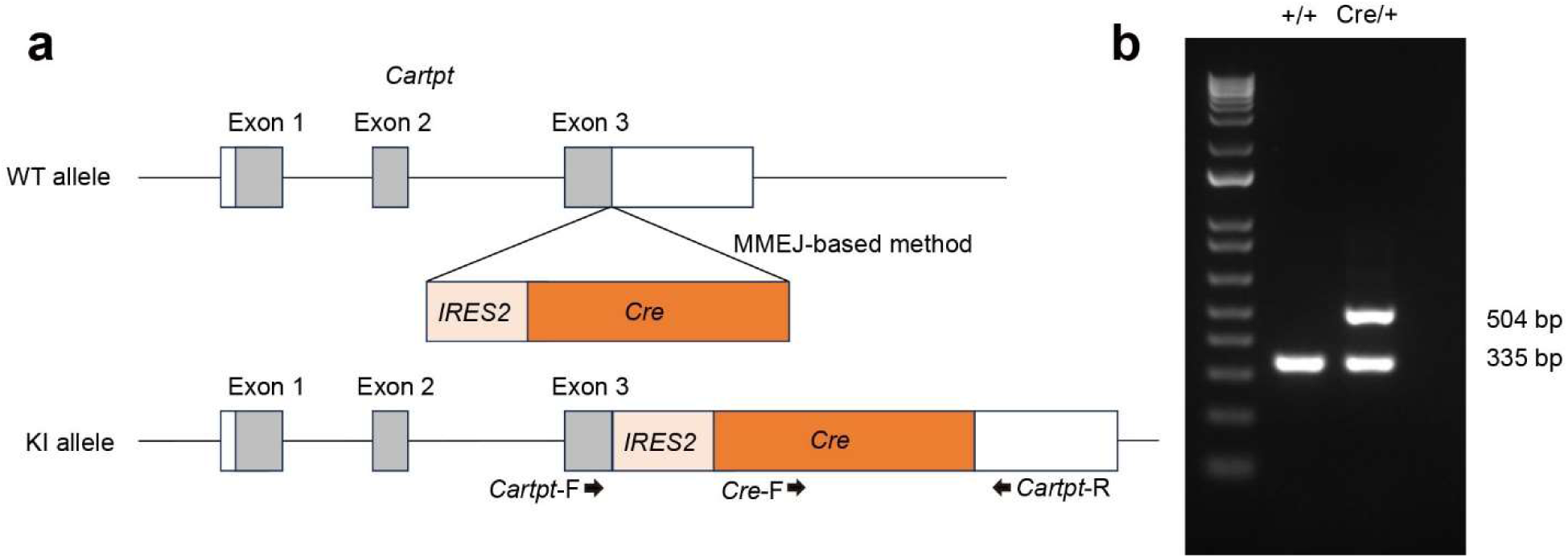
Generation of *Cartpt-Cre* mice, related to Fig. 2. **a**, Illustration of the knock-in strategy for generating *Cartpt-Cre* mice using the microhomology-mediated end joining (MMEJ)-based method^26^. Top: The *Cartpt* gene comprises three exons, where open and closed boxes denote untranslated and protein-coding regions, respectively. The *IRES2-Cre* cassette was inserted immediately downstream of the coding end of *Cartpt* on exon 3. Bottom: Schematic representation of the knock-in allele. Black arrows indicate the location of genotype primers. **b**, Representative gel image displaying electrophoresis results examining the *Cre* allele and negative control (wild-type animal).

**Extended Data Fig. 3:**
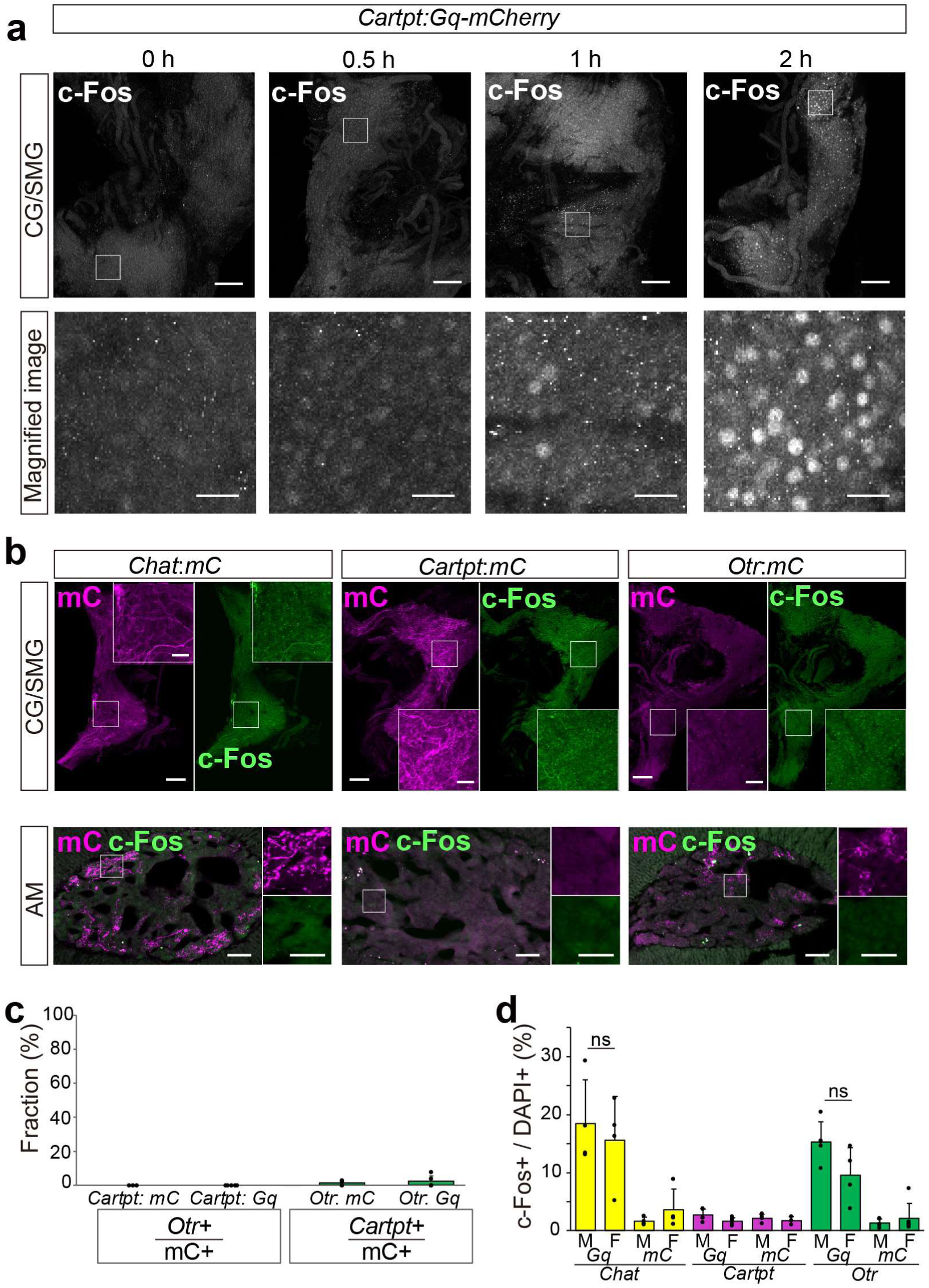
Time course and control experiments for the activation of specific SPNs, related to Fig. 2. **a**, Expression of c-Fos (gray) in the whole mount view of the CG/SMG is shown at various time points (0, 0.5, 1, or 2 h) following CNO administration in *Cartpt-Cre* mice injected with AAV8-*hSyn-DIO-Gq-mCherry* into the T8–T12 SC. Lower panels display magnified images within the white-boxed areas. Scale bars, 200 μm for the top images and 50 μm for the magnified images. **b**, Expression of c-Fos (green) in the whole mount view of the CG/SMG (upper) and adrenal medulla (AM) sections (lower) of *Chat-Cre*, *Cartpt-Cre*, or *Otr-Cre* mice injected with AAV8-*hSyn-DIO-mC* (for control) in the SC, observed 2 hours after CNO administration. mC+ axons are shown in magenta. Scale bars, 200 μm for low-magnification images and 50 μm for magnified images. **c**, Percentage of mC+ cells in *Cartpt-Cre* mice that express *Otr* mRNA (left) and in *Otr-Cre* mice that express *Cartpt* mRNA (right). The expression of mC was induced using AAV8-*hSyn-DIO-mCherry* and AAV8-*hSyn-DIO-Gq-mCherry*. N = 3–5 mice. These data support the high specificity of Cre/AAV-mediated targeting within specific SPNs. **d**, The fraction of c-Fos+ per DAPI+ cells in the AM of CNO-treated *Gq* and *mC* control male (M) and female (F) mice. Data correspond to Fig. 2g; however, data are presented separately for males and females. N = 4–6. Data are presented as mean ± standard deviation. Statistical comparisons were conducted using a two-sided Welch’s *t*-test (**d**).

**Extended Data Fig. 4:**
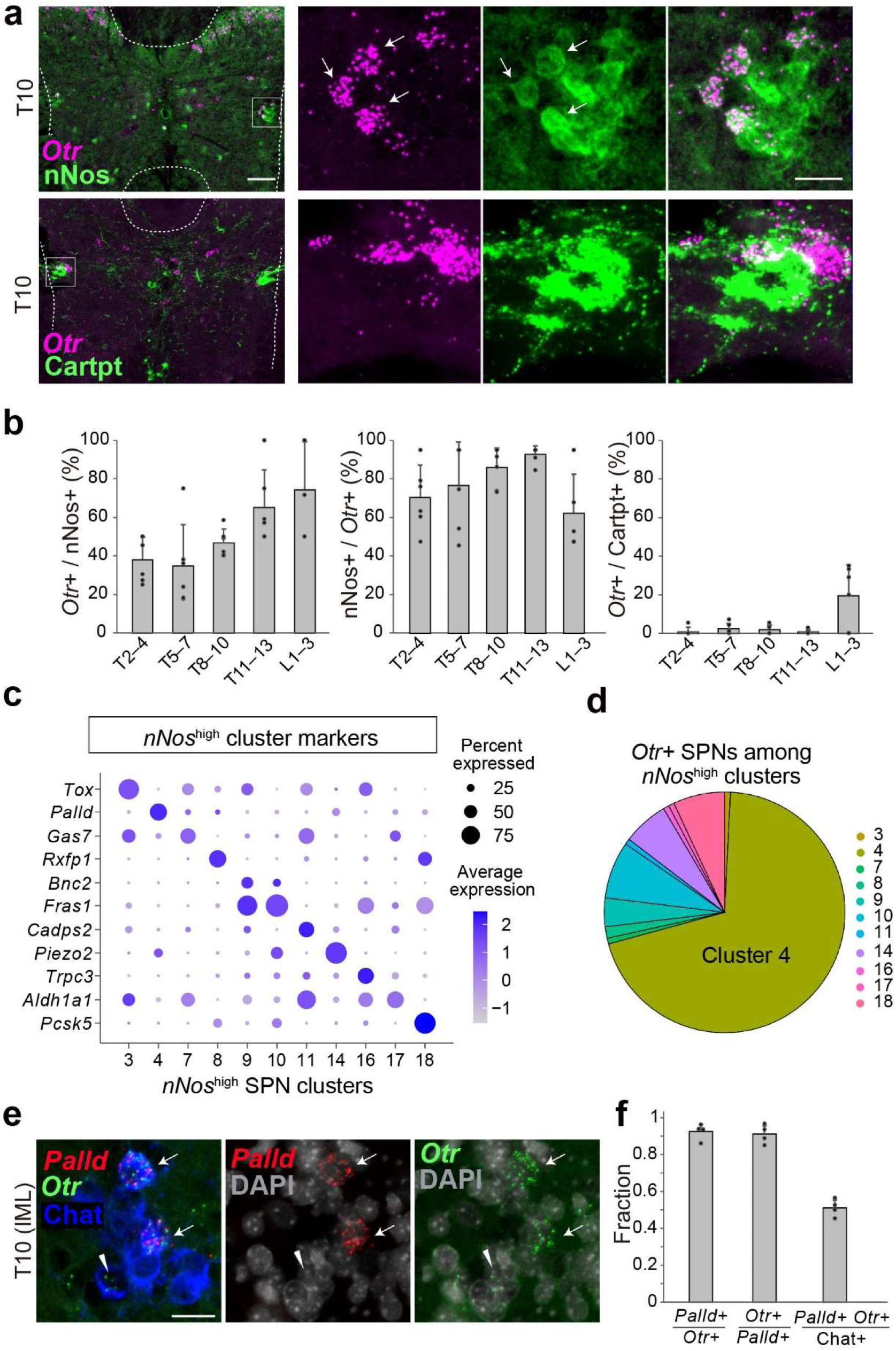
*Otr* marks a *nNos*+ TC type corresponding to cluster 4, related to Fig. 2. **a**, Expression of *Otr* mRNA (detected by RNAscope, magenta) and nNos or Cartpt immunostainings (green) in the T10 SC of wild-type mice. The right panels present magnified and channel-separated images within the white-boxed area. Arrows indicate cells that are dual-positive for *Otr* and nNos. Scale bars, 100 μm for the left images and 25 μm for the magnified images. **b**, Percentage of *Otr*+ cells among nNos+ cells (left), nNos+ cells among *Otr*+ cells (middle) in the IML regions, and *Otr*+ cells among Cartpt+ cells (right) in the IC, CA, and IML regions. N = 6 mice. **c**, Identification of enriched differentially expressed genes among SPNs for *nNos*^high^ clusters, as in Fig. 1i. **d**, Pie chart displaying the fraction of *Otr*+ SPNs detected within each *nNos*^high^ SPN cluster. The majority of *Otr*+ SPNs are found in cluster 4. **e**, Expression of *Palld* (red, a cluster 4 marker) and *Otr* (green) mRNAs detected by RNAscope, along with Chat immunostaining (blue) and DAPI staining in the IML of the T10 spinal segment of wild-type mice. Arrows indicate cells that are triple-positive for *Palld*, *Otr*, and Chat, while the arrowhead depicts a *Palld*– *Otr*+ Chat+ cell. Scale bars, 25 μm. **f**, Percentage of *Palld*+ cells among *Otr*+ cells (left), *Otr*+ cells among *Palld*+ cells (middle), and *Palld+ Otr*+ dual-positive cells among Chat+ cells (right) in the IML of the T8–T13 segments. N = 4 mice. Data are presented as mean ± standard deviation.

**Extended Data Fig. 5:**
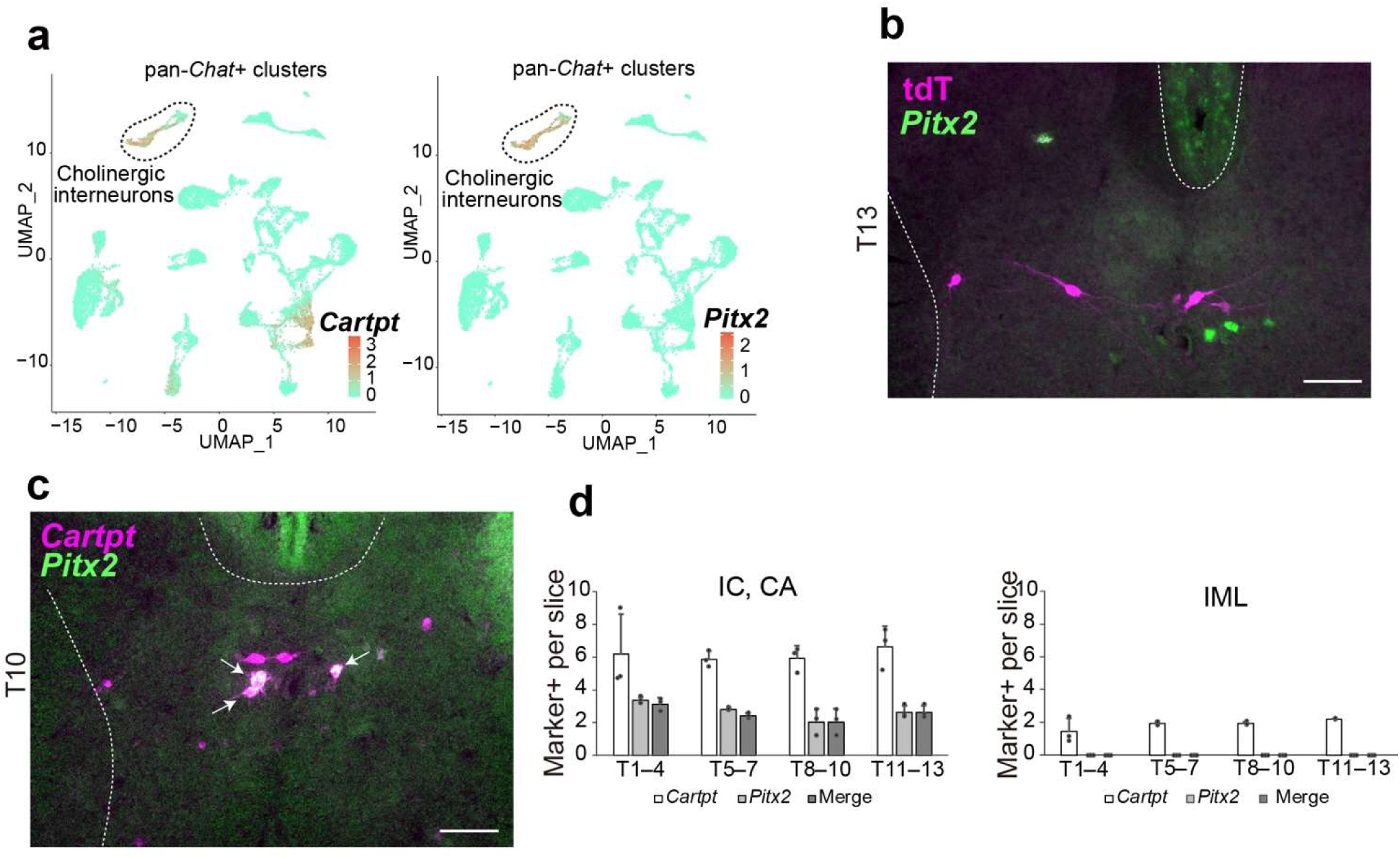
*Cartpt*+ cholinergic interneurons and projection-specific manipulation of CG/SMG-projecting SPNs, related to Fig. 3. **a**, UMAP representation of pan-*Chat*+ clusters based on the reanalysis of published transcriptome data^12^ with a colored heat map showing the log-normalized expression of *Cartpt* (left) and *Paired-like homeodomain 2* (*Pitx2*), a marker of the cholinergic interneurons (right). The dashed line-enclosed clusters represent *Pitx2*+ cholinergic interneurons, which are also *Cartpt+.* **b**, Coronal section of the T13 SC displaying retrogradely labeled tdT+ cells (magenta) from the CG/SMG and *Pitx2* mRNA (green) expression. Scale bars, 100 μm. No overlap was observed. **c**, Expression of *Cartpt* mRNA (magenta) and *Pitx2* mRNA (green) in the T10 SC of wild-type mice. The arrows indicate cells that are double-positive for *Cartpt* and *Pitx2*. Scale bars, 100 μm. **d**, Average number of *Cartpt*+ (white), *Pitx2*+ (light gray), and *Cartpt*+ *Pitx2*+ (dark gray) cells per slice in the IC and CA (left) or IML (right) of the T1–4, T5–7, T8–10, and T11–13 SC segments in wild-type mice. N = 3.

**Extended Data Fig. 6:**
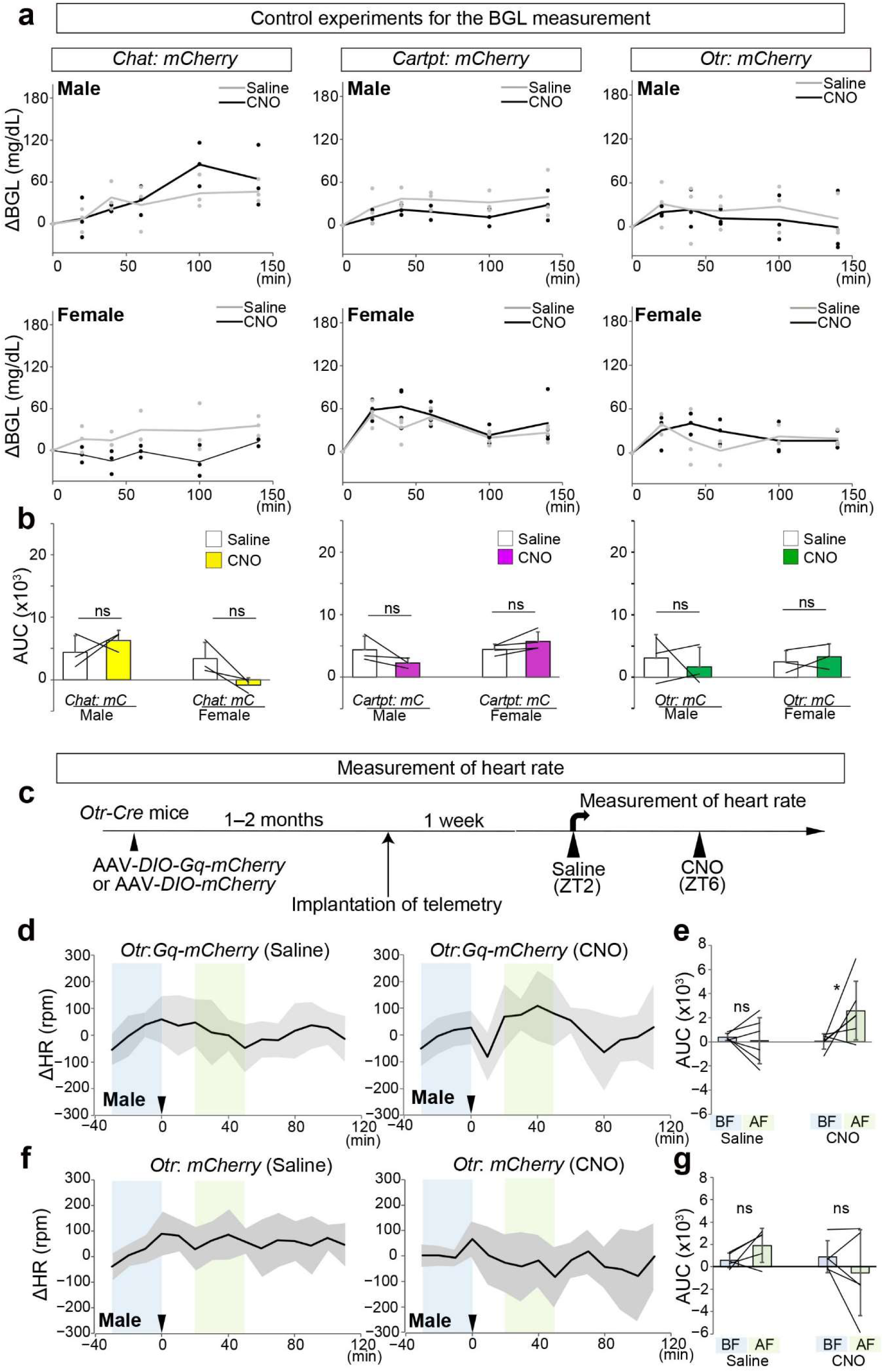
Control data on blood glucose levels and measurement of heart rate, related to Fig. 4. **a**, ΔBGL in *Chat-Cre*, *Cartpt-Cre*, or *Otr-Cre* mice with AAV8-*hSyn-DIO-mCherry* injection (for control) upon administration of saline (gray) or CNO (black). Individual results are represented by dots, while the average is indicated by a line. The upper and lower graphs depict results for male and female mice, respectively. **b**, AUC measurements for 0 to 140 minutes. N = 3–4 mice per condition. **c**, Schematics illustrating the measurement of heart rate using a telemetry system. ZT, zeitgeber time, with ZT0 representing 08:00. **d**, **f**, Change in heart rate (ΔHR) in *Otr-Cre* male mice injected with AAV8-*hSyn-DIO-Gq-mCherry* (panel **d**) or AAV8-*hSyn-DIO-mCherry* injection (for control, panel **f**) into the T8–T12 spinal segments, upon administration of saline (left) or CNO (right). The average is indicated by a line, with gray shadows representing the standard deviation. Arrowheads indicate the timing of saline or CNO administration. Light blue and light green areas indicate –30 to 0 minutes and +20 to +50 minutes relative to the saline or CNO administration, respectively. **e**, **g**, AUC measurements for –30 to 0 minutes and +20 to +50 minutes. N = 5–6 mice. Data are presented as mean ± standard deviation. Statistical comparisons were made using two-way ANOVA with repeated measures; (*Chat-Cre*) sex effect, *p* < 0.05, solution effect, ns, interaction effect, ns. (*Cartpt-Cre*) sex effect, ns, solution effect, ns, interaction effect, ns. (*Otr-Cre*) sex effect, ns, solution effect, ns, interaction effect, ns (**b**). Two-way ANOVA with repeated measures; (*Otr*: *Gq, Otr*: *mC*) interaction effect, ns, \**p* < 0.05 by a post hoc two-sided Welch’s *t*-test with Bonferroni correction (**e, g**).

**Extended Data Fig. 7:**
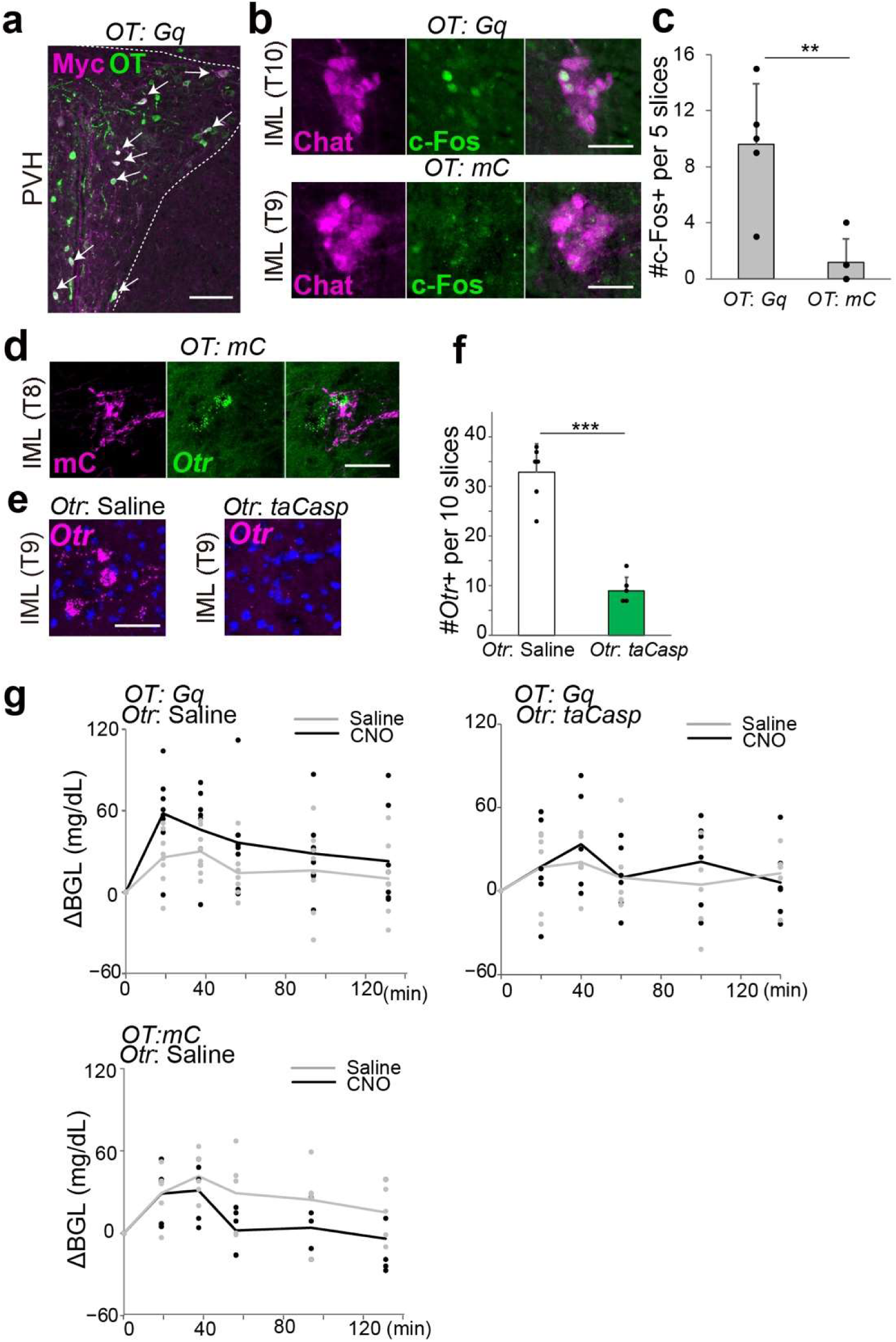
Additional data on ablation of *Otr*+ SPNs, related to Fig. 4. **a**, Coronal section of the posterior showing the expression of Myc (magenta) fused with Gq and OT (green) following the injection of AAV5-*OTp*-*Gq-Myc*. Arrows indicate cells that are double-positive for Myc and OT. This area corresponds to the posterior part of the PVH where parvocellular PVH OT neurons projecting the SC present^65^. **b**, Coronal section of the spinal cord segment showing the expression of Chat (magenta) and c-Fos (green) in IML region following CNO administration in *Otr-Cre* mice injected with AAV5-*OTp*-*Gq-Myc* or AV9-*OTp*-*mCherry* into the PVH. **c**, Number of c-Fos+ cells in 5 sections of the T8–12 IML. N = 5 each. **d**, Coronal section of the T8 spinal segment showing the expression of mC (magenta) and *Otr* (green) following the injection of AAV9-*OTp*-*mCherry*. This data demonstrates direct axonal projections from PVH OT neurons to the IML of lower thoracic SC. **e**, Coronal section of the T9 spinal segment in *Otr-Cre* mice injected with saline control (left) or AAV1-*FLEx-taCasp3-TEVP* (right). *Otr* mRNA expression is depicted by RNAscope in magenta and DAPI staining in blue. Scale bars, 100 μm (panels **a**, **b**, **d**, **e**). **f**, Number of *Otr*+ cells in the control saline-injected group (white) and the ablation group (green) across 10 sections from the T8–12 SC. N = 6 each. **g**, ΔBGL in the OT neuron-activated group (top left), OT neuron-activated and *Otr+* SPN-ablated group (top right), and control group (bottom) upon administration of saline (gray) or CNO (black). Baseline glucose levels at time 0 minutes did not differ: 172.13 ± 14.89 mg/dL for *OT*: *Gq*, *Otr*: Saline, 165.75 ± 17.68 mg/dL for *OT*: *Gq*, *Otr*: *taCasp*, and 150.40 ± 13.14 mg/dL for control. N = 5–8. Data are presented as mean ± standard deviation. **, *p* < 0.01, ***, *p* < 0.001 by a two-sided Welch’s *t*-test (**c**, **f**).

**Supplementary Movie 1: Light-sheet microscopy of the CG/SMG, related to Fig. 1.**

Whole mount view of the CG/SMG of double heterozygous *DBH-CreER^T2^*; *Ai9* mice with tamoxifen administration, followed by CUBIC-based tissue clearing. Magenta cells represent postGNs labeled with tdT from the *Ai9* allele.

**Supplementary Movie 2: Light-sheet microscopy of the SC, related to Fig. 1.**

Whole mount view of the tissue-cleared T5–13 SC of *Ai9* mice that received AAVrg *hSyn-Cre* injection into the CG/SMG, displaying retrogradely labeled tdT+ cells (magenta).

## Notes

### Competing Interest Statement

The authors have declared no competing interest.

